# Contribution of cytotoxic CD8 T cells, neutrophils and type 1 interferon signaling to hyperinflammation in HIV-associated TB meningitis

**DOI:** 10.64898/2026.03.10.710544

**Authors:** James R Barnacle, Nonzwakazi Bangani, Hubert Slawinski, Christopher Barrington, Katalin A Wilkinson, Cari J Stek, Rachel Lai, Fabrice Bonnet, Graeme Meintjes, Brian Robertson, Martin Gengenbacher, Angharad G Davis, Daniel L Barber, Anne O’Garra, Robert J Wilkinson

**Affiliations:** Francis Crick Institute, London, NW1 1AT, UK; Department of Infectious Diseases, Imperial College London, W12 0NN, UK; Wellcome Discovery Research Platforms in Infection, Centre for Infectious Diseases Research in Africa, Institute of Infectious Disease and Molecular Medicine, University of Cape Town, Observatory 7925, Republic of South Africa; Genomics Science Technology Platform, Francis Crick Institute, London, NW1 1AT, UK; Bioinformatics and Biostatistics Science Technology Platform, Francis Crick Institute, London, NW1 1AT, UK; University of Bordeaux, National Institute for Health and Medical Research (INSERM) UMR 1219, Research Institute for Sustainable Development (IRD) EMR 271, Bordeaux Population Health Centre, Bordeaux, France; Centre for Immunobiology and Infection, Blizard Institute, Queen Mary, University of London, London, E1 2AT, UK; Department of Medicine, Faculty of Health Sciences, University of Cape Town, Cape Town, Observatory 7925, Republic of South Africa; Center for Discovery and Innovation, NJ 07110, USA; T Lymphocyte Biology Section, Laboratory of Parasitic Diseases, National Institute of Allergy and Infectious Diseases, National Institutes of Health, Bethesda, MD, 20892, USA; Immunoregulation and Infection Laboratory, Francis Crick Institute, London, NW1 1AT, UK

## Abstract

Immune dysregulation contributes to death and disability in tuberculous meningitis. People living with HIV have the least evidence that anti-inflammatory therapy improves the poor outcome. Improving therapy relies on a more refined understanding of the host immune response. Single-cell RNA sequencing of 188,983 CSF cells from 25 adults with HIV-associated TBM revealed a predominance of cytotoxic CD8 T cells with low cytokine expression. In microbiologically-confirmed TBM, there was greater cytotoxicity in T, NK and γδ cells, and higher type 1 interferon stimulation in T and B lymphocytes. Neutrophils expressed markers suggesting heightened cytokine stimulation, enhanced effector function, and IL-8-mediated neutrophil recruitment. In a longitudinal cohort, type 1 interferon signaling increased in blood and CSF following treatment initiation. Overall, findings indicate a hyper-inflammatory immune response in the CSF of HIV-associated TBM patients characterised by an accumulation of granzyme-rich cytotoxic CD8 T cells, highly activated neutrophils and host-detrimental type 1 interferon signaling.

## INTRODUCTION

Tuberculosis (TB) is the leading infectious cause of death globally with an estimated 10.8 million people falling ill and 1.25 million deaths from TB in 2023 (*1*). Tuberculous meningitis (TBM) is the most severe form of TB. In 2019, there were an estimated 164,000 adult cases worldwide causing 78,200 deaths (*2*). Although the global prevalence of human immunodeficiency virus type 1 (HIV-1) is ∼0.5%, 23% of TBM cases and 35% of deaths are in people living with HIV/AIDS (PLWHA) (*2*). In HIV-uninfected patients the 6-month mortality from TBM is 24%, with physical disability reported in 32% of adult survivors. In PLWHA, mortality can be as high as 57% (*3*). Recent studies have found no benefit (and even a suggestion of harm) from higher doses of rifampicin (the cornerstone antibiotic in TB therapy) suggesting that improving outcomes cannot solely be mediated by rapid bacterial killing (*4*, *5*). Immune dysregulation contributes to poor outcome, and host-directed adjunctive therapies may modulate this immunopathology. However the biggest clinical trial in HIV-TBM showed corticosteroids, hitherto the mainstay of adjunctive therapy for TBM, did not confer benefit with respect to survival or any secondary end point (*6*). Therefore, PLWHA are at the greatest risk of developing and dying from TBM but appear to benefit least from existing adjunctive therapies. Developing new therapies is hindered by a lack of understanding of the central nervous system (CNS) immune response to TBM.

Like other forms of HIV-associated extrapulmonary TB, the bacterial load in TBM varies widely between patients (*7*, *8*). This is most often ascribed to diagnostic insensitivity but may also reflect unsuspected differences in host-pathogen interaction. Leprosy, another mycobacterial disease, presents as a spectrum of disease ranging between paucibacillary tuberculoid and multibacillary lepromatous disease which have distinct host transcriptomic correlates, suggesting an intimate relationship between antimycobacterial immunity in humans and bacterial load (*9*). In TBM, higher CSF bacterial loads have been associated with disease severity, CSF neutrophilia, cytokine concentrations, neurological events, and mortality (*7*, *10*).

Single cell RNA sequencing (scRNAseq) provides high-resolution phenotyping of immune cells. Thus far, its use in TBM has been restricted to three small studies not powered to encompass patient and disease heterogeneity or make comparisons between groups. One study identified *Tnfrsf4/OX40* expression as a possible marker of recently activated CD4 T cells in mice and demonstrated *OX40* expression in sorted CD4 T cells in the CSF of a single PLWHA with TBM (*11*). Another sequenced CSF and blood cells from six children with TBM and identified a complement activated microglia-like population unique to CSF (*12*). A third study sequenced CSF and blood cells from four HIV-uninfected adult males with TBM, of whom two were microbiologically positive and one of whom died within a week of admission. CSF was enriched with highly inflammatory microglia-like macrophages, GZMK+CD8+ effector-memory T (TEM) cells, and CD56bright NK cells compared to blood (*13*).

We hypothesised that TBM patients with higher and lower bacterial loads would display distinct immune responses. Using scRNAseq, this study aimed to describe the CSF immune cell composition and phenotype in patients with HIV-associated TBM and compare the immune response in patients with and without detectable bacilli in the CSF.

## RESULTS

### Single cell RNA sequencing showed heterogenous immune cell composition in the lumbar CSF in HIV-associated TBM but with consistent CD8 T cell predominance

ScRNAseq was performed on 25 lumbar CSF samples from South African PLWHA aged ≥15 years with a diagnosis of possible, probable or microbiologically positive TBM using 10x Genomics Fixed RNA Profiling (**Fig. 1A**). Participants were enrolled in the INTENSE-TBM trial and had a lumbar puncture seven days following enrolment. Study demographics and individual patient clinical data can be found in **Table S1 and S2**, respectively.

**Fig. 1.**
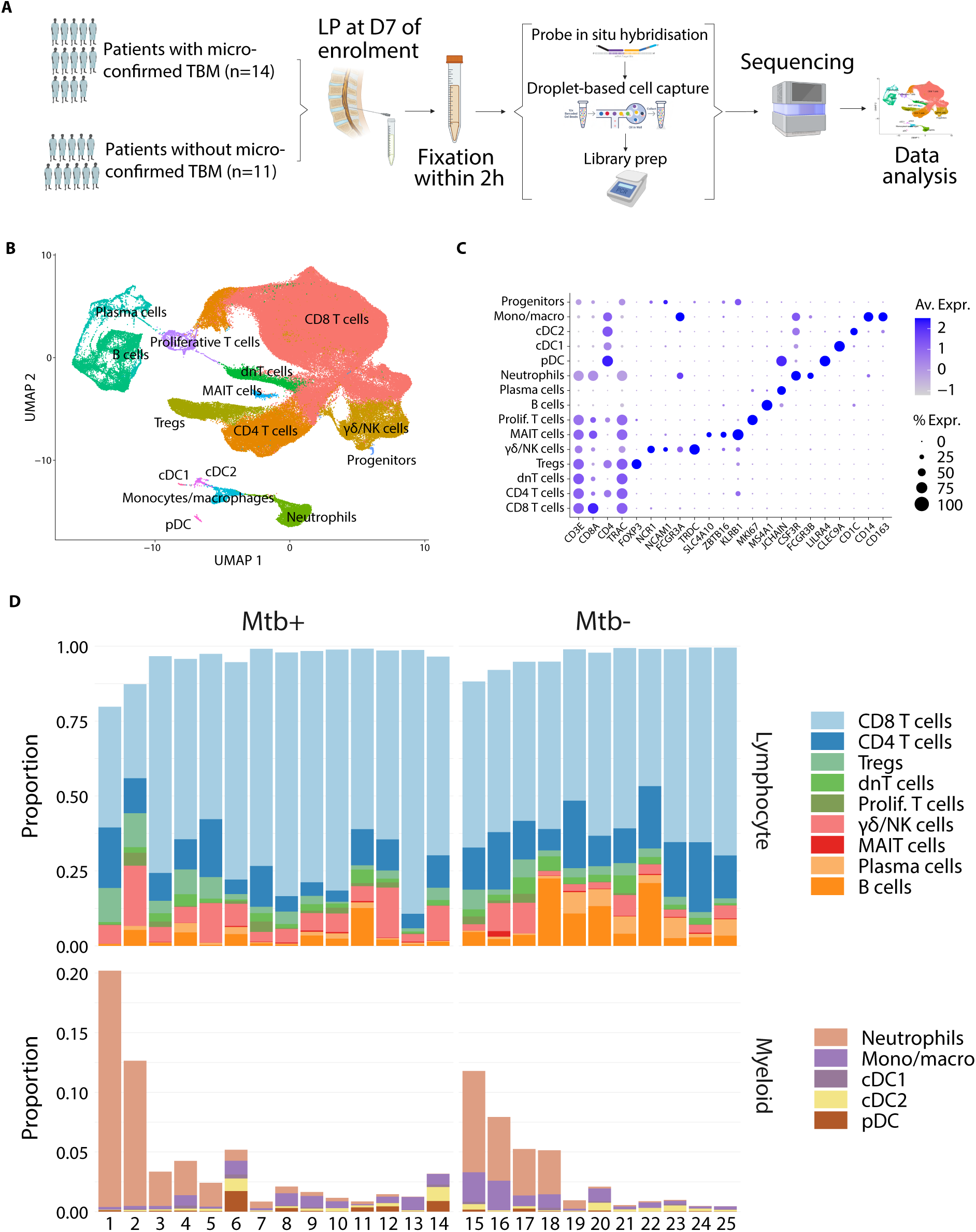
CD8 T cells predominate in the CSF in HIV-associated TBM. **(A)** Study design. **(B)** Annotated UMAP of 188,983 cells from the lumbar CSF of 25 adults with HIV-associated TBM. **(C)** Dot plot of canonical gene expression of annotated clusters. **(D)** Cell proportions of annotated clusters by microbiological confirmation. Split by myeloid and lymphocyte lineages and ordered by neutrophil proportion within each microbiological status. Note that myeloid y axis has been cut at 0.2 to aid visualisation. ART, anti-retroviral therapy; pDC, plasmacytoid dendritic cells; cDC, conventional dendritic cell; NK, natural killer; MAIT, mucosal-associated invariant T cells; Tregs, regulatory T cells; dnT, double-negative T cells; γδ, gamma-delta.

A total of 188,983 cells were identified from all samples after ambient RNA contamination and doublet removal (**B**). Coarse cell type annotation was done using both canonical marker genes and differentially expressed genes between clusters, checked against automated annotation methods (**C**). Cell type proportions revealed significant heterogeneity between patients (**D**). CD8 T cells were the predominant cell type (63.1% [patient range 31.4-88.0%]), followed by CD4 T cells (12.2% [3.8-23.4%]), γδ/NK cells (6.1% [1.6-20.2%]), B cells (5.4% [0.5-22.5%]), neutrophils (2.4% [0-19.8%]), double-negative T cells (2.3% [0.6-6.0%]), plasma cells (2.0% [0.08-7.1%]), proliferating T cells (1.1% [0.03-4.3%]), monocytes/macrophages (0.8% [0.09-2.5%]), conventional dendritic cells (cDC2) (0.3% [0.07-1.2%]), mucosal-associated invariant T (MAIT) cells (0.3% [0-1.9%]) and plasmacytoid dendritic cells (pDC) (0.2% [0-1.7%]), cDC1 (0.1% [0-0.4%]) (**Table S4**). All participants had higher proportions of CD8 T cells than CD4 T cells, even those participants with CD4 counts >500 cells/mm^3^. Plasma cells and B cells showed a strong positive correlation as expected. CD8 T cells correlated positively with non-neutrophil myeloid cells, whereas regulatory T cells (Tregs) and neutrophils co-correlated, but had a strong negative correlation with MAIT cells (**Fig. S2**).

### The majority of CD8 T cells in the CSF were highly cytotoxic with limited cytokine expression

To investigate phenotypic heterogeneity within broad cell annotations, CD8 T cells, CD4 T cells, NK and γδ cells, B and plasma cells, and myeloid cells were individually subsetted, re-clustered and re-annotated (**Fig. S3 to S7**). The majority of CD8 T cells (n = 123,557) had a cytotoxic effector memory-like phenotype with high natural killer cell granule protein 7 (*NKG7*), granzyme A, M, K (*GZMA*, *GZMM*, *GZMK*) and perforin (*PRF1*) expression (**Fig. 2A**). Amongst these cytotoxic CD8 T cells, there were populations with high granulysin (*GNLY*) expression and an NK-like phenotype expressing natural cytotoxicity triggering receptor 1 (*NCR1*) and killer cell lectin like receptor (*KLRC1*, *KLRC3*, *KLRC4*) (**B).** Cytokine expression in CD8 T cells was low. Th1/Th17-associated genes, including *IFNG*, were mostly limited to a single cluster, which co-expressed *IL23R*, *VDR*, *CD226*, and *RORC*. This IFNG+ cluster also expressed granzyme B (*GZMB*), which was only seen elsewhere in the GNLY+ cluster (B). Both of these GZMB+ clusters were significantly increased in microbiologically positive disease in composition analysis (**C, Fig. S3E**). Amongst CD4 T cells (n = 21,844), there were high numbers of FOXP3+ regulatory T cells and cells expressing naïve and central memory (*IL7R*, *CCR7*, *TCF7*, *SELL*, *LEF1*) (**D**). As with CD8 T cells, cytokine expression overall was low, but an IFNG+ cluster was again seen, which expressed cytotoxicity markers, in particular *PRF1, GZMA* and *GZMB* (**E**). This IFNG+ cluster was not increased in microbiologically positive disease, unlike in the CD8 T cell compartment (**F, Fig. S4E**). NK and γδ cells (n = 11,483) did not separate clearly on the main UMAP, suggesting overlap in function and gene expression, and were subsetted and re-clustered together (**G**). *NCAM1* (CD56) and *FCGR3A* (CD16) expression was inversely correlated in NK cells (**H**). A small γδ cluster had a naïve-like phenotype with high *LEF1*, *TCF7* and *IL7R* expression. The other two γδ clusters had cytotoxic phenotypes with high granzyme, perforin and *NKG7* expression, with one cluster also having high *IL7R*, *CXCR6*, *KLRB1* and *KLRG1* expression, and the other with high *TIGIT* expression (H).

**Fig. 2.**
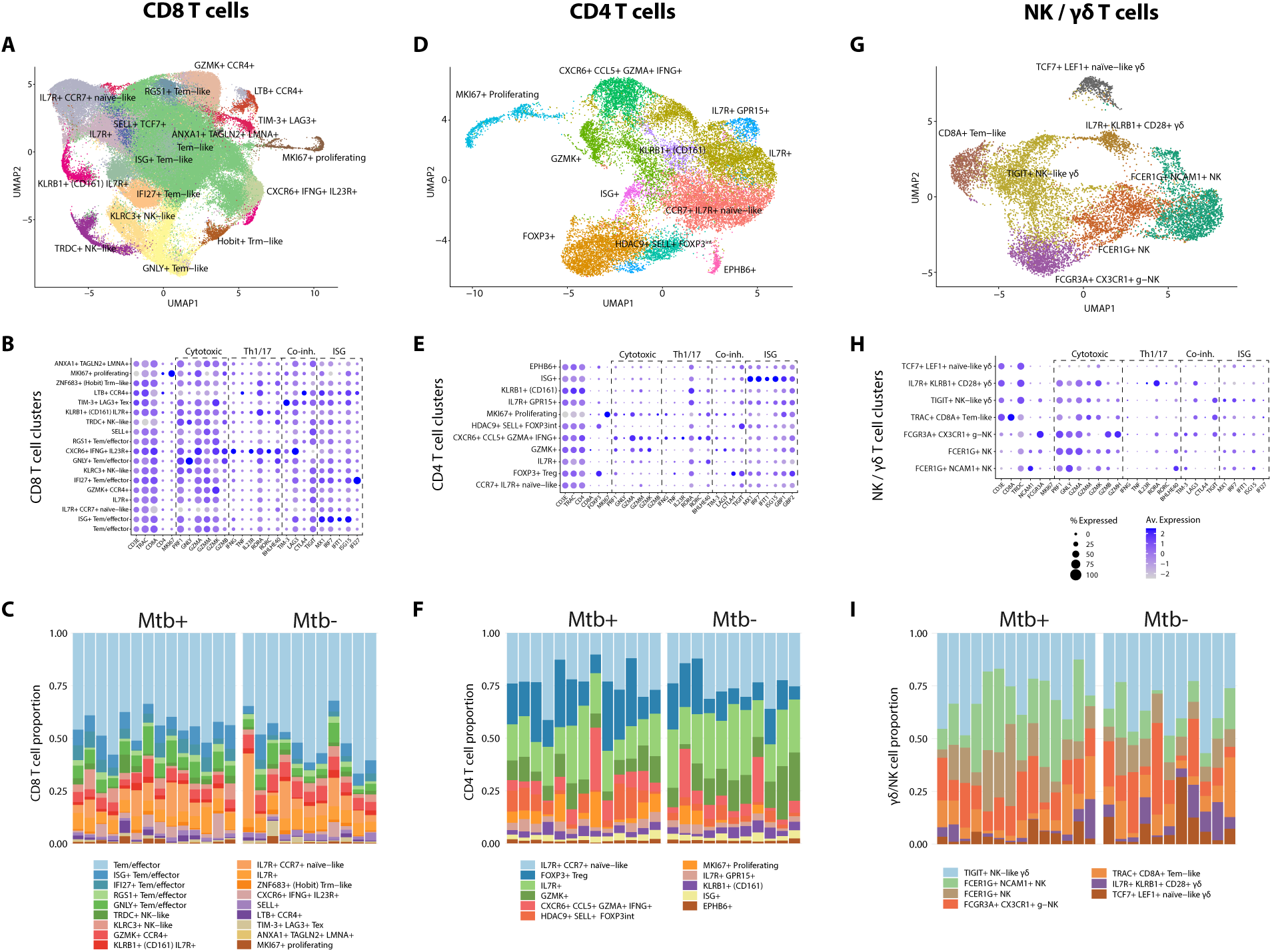
Phenotypic features of CD8, CD4, γ8 and NK cells. **(A)** Annotated UMAP of 119,825 CD8 T cells. **(B)** Dotplot of CD8 T cell clusters showing expression of genes associated with cytotoxicity, interferon stimulation, Th1/Th17 and inhibitory co-receptor expression. **(C)** Cell proportions of annotated CD8 T cell clusters by microbiological confirmation. **(D)** Annotated UMAP of 21,844 CD4 T cells. **(E)** Dotplot of CD4 T cell clusters showing expression of genes associated with cytotoxicity, interferon stimulation Th1/Th17 and inhibitory co-receptor expression. **(F)** Cell proportions of annotated CD4 T cell clusters by microbiological confirmation. **(G)** Annotated UMAP of 11,483 NK / γ8 cells. **(H)** Dotplot of NK / γ8 cell clusters showing expression of genes associated with cytotoxicity, interferon stimulation Th1/Th17 and inhibitory co-receptor expression. **(I)** Cell proportions of annotated NK / γ8 cell clusters by microbiological confirmation. *Mtb+,* microbiologically confirmed TBM.

B cell and plasma cells (n = 14,042) separated into distinct naïve B cell, memory B cell, plasma cell and plasmablast clusters. Memory B cell clusters were a mix of class-switched and non-class-switched B cells (**Fig. S6**). Myeloid cells (n = 7,182) from the neutrophil, monocyte/macrophage and dendritic cell clusters were subsetted and re-clustered (**Fig. S7A to S7D**). Of the nine clusters, four were neutrophils (clusters 0, 2, 4, 6), two were pDC (clusters 3, 8), one was cDC1, one was macrophages and the remaining cluster was a mix most likely containing monocytes, monocyte-derived macrophages (MDM), microglia, and CNS-associated macrophages (CAM), including perivascular, meningeal and choroid plexus macrophages (*14*).

Overall, the lumbar CSF of patients with HIV-associated TBM showed a granzyme-rich cytotoxic CD8 T cells predominance, even in patients with CD4 counts >500 cells/mm^3^. However, cell type proportions differed considerably between patients and, in particular, proportions of IFNG+ CD8 T cells were higher in patients with microbiologically positive disease.

### Cytotoxic mediators were overexpressed in CD4, MAIT and CD56^hi^ NK cells in microbiologically positive TBM

**Figure 3A** shows cytotoxic gene expression across all cells. To compare differences between patients in cell type and transcript abundance by microbiological status at enrolment, pseudobulk differential gene expression (DEG) and composition analysis were performed. CD4, pDC, Treg, MAIT and γδ/NK cells had the highest numbers of DEG between patients with and without microbiologically confirmed TBM (**C**). Patients with higher bacterial loads had increased expression of cytotoxic genes (*GZMB*, *GZMH*, *PRF1*, *NKG7*) in their CSF, in particular CD4 T cell and MAIT cells (C). When DEG was done on cell subclusters, amongst CD8 T cell, the Tem/effector and GNLY+ Tem/effector clusters significantly overexpressed GNLY and the NK cell marker natural cytotoxicity triggering receptor 1 (*NCR1*), respectively, in microbiologically confirmed disease (**D**). In CD4 T cell subclusters, *GZMB* expression in the IFNG+ cluster was almost 25X higher in microbiologically positive TBM (**Fig. S4F**). Functional enrichment analysis of DEG in this IFNG+ CD4 T cell cluster showed upregulation of oxidative phosphorylation, cell killing and nucleic acid synthesis, suggesting cells in this cluster exhibit increased metabolism and activation in patients with higher *M. tuberculosis* bacterial loads in CSF (**Fig. S4G**). Amongst NK subclusters, *GZMB*, *GNLY* and *KLRG1* expression was significantly increased in FCER1G+ (Fc epsilon receptor subunit gamma) clusters in microbiologically confirmed TBM (D).

**Fig. 3.**
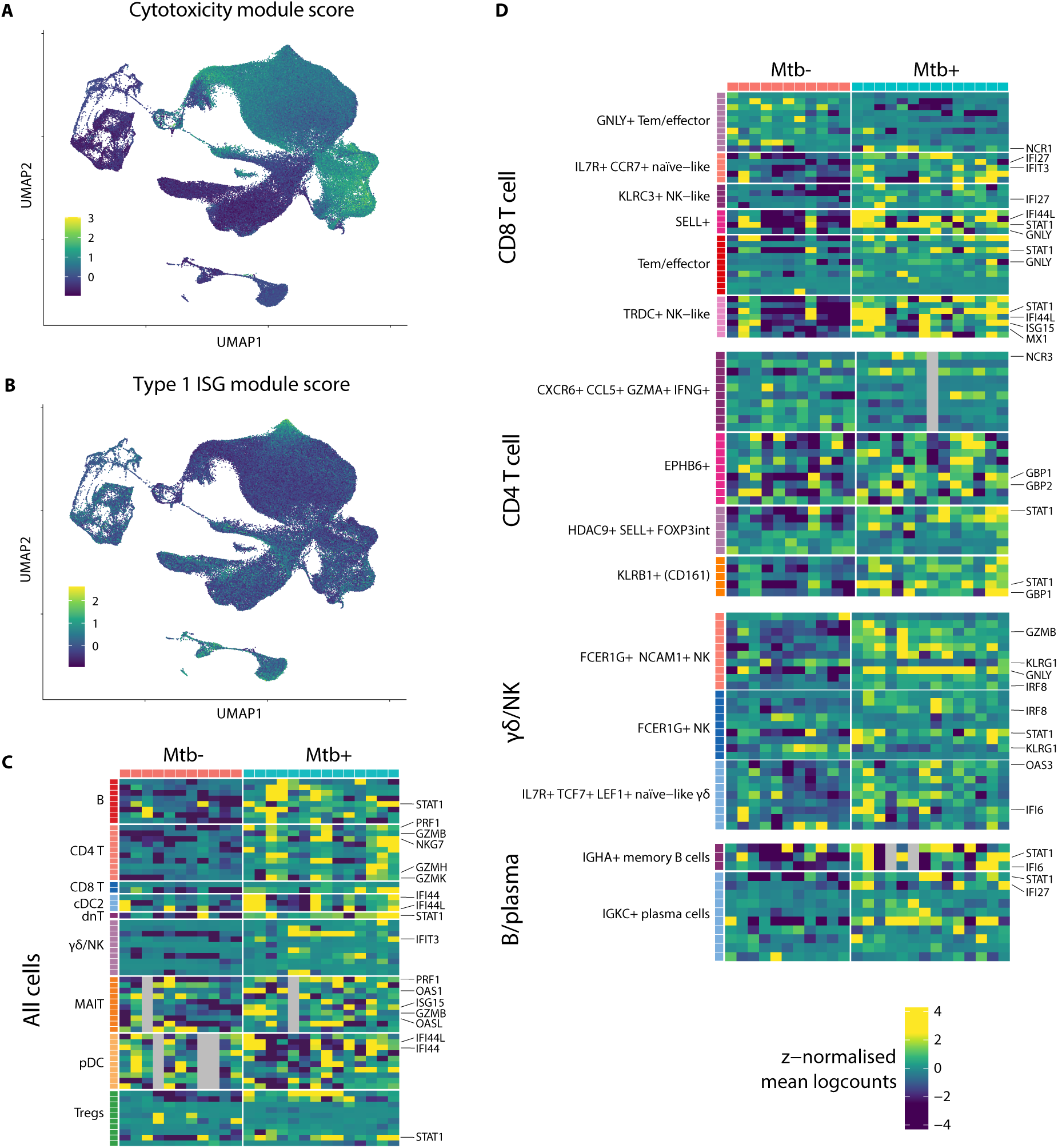
Genes associated with cytotoxicity and type 1 interferon stimulation are overexpressed in microbiologically confirmed disease. **(A)** UMAP of cytotoxicity gene module expression in all cells. **(B)** UMAP of type 1 interferon associated gene module expression in all cells. **(C)** DEG associated with cytotoxicity and type 1 interferon stimulation in each main cell type cluster between those with and without microbiological confirmation. Pseudobulk differential expression analysis was calculated using the muscat R package with DESeq (Wald test) with age and sex as model co-variates. If fewer than 10 DEG then all shown. Rows split by cluster, columns by microbiological status. Grey shading indicates that the patient had too few cells in that cluster to calculate. **(D)** DEG in each cell subcluster between those with and without microbiological confirmation. Only subclusters containing significant genes are included. Tem, effector memory T cell; Trm, tissue resident memory T cells; *KLRB1*, Killer cell lectin-like receptor subfamily B, member 1 (CD161); NK, natural killer; *GNLY*, granulysin; *IFNG*, interferon gamma; ISG, interferon-stimulated gene; *MKI67*, Kiel 67 (Ki-67).

Proliferating T cells, pDC and γδ/NK cells were significantly more abundant in patients with microbiologically positive TBM, whereas plasma and B cells were significantly lower (**Fig. S1F**). Interestingly, when comparing the cell compositions of the eight patients who were on ART at enrolment with patients not on HIV treatment, patients on ART had very few proliferating T cells in their CSF. Given that the HIV-1 viral load was higher in patient with microbiologically confirmed TBM, these increased proliferating cells may be responding to HIV-1 antigen, rather than *M. tuberculosis*. Intuitively, CD4 T cell proportions were significantly higher in those on ART treatment along with MAIT cells, whilst cDC1 and pDC were decreased (**Fig. S1G**).

Overall, cytotoxic mediators were overexpressed in CD4, NK and MAIT cells in microbiologically positive TBM, suggesting that the cytotoxicity seen in the lumbar CSF in HIV-associated TBM is further increased in patients with higher bacterial loads.

### Neutrophils were highly activated and separated into two phenotypes expressing increased IL-8 and CXCR2 respectively

4,606 neutrophils were identified after re-clustering and annotating the 7,182 myeloid cells (**Fig. 4A to 4B**). These neutrophils globally expressed colony-stimulating factor 3 receptor (*CSF3R*), intercellular adhesion molecule 1 (*ICAM1*), and nicotinamide phosphoribosyltransferase (NAMPT, visfatin) (**C**). Neutrophils either had higher *MMP25*, *IL1B* and *SOCS3* (suppressor of cytokine signaling 3) expression, or higher *CXCL8* (IL-8) expression in association with either *CXCR4* or *NFXL1* (nuclear transcription factor, X-box binding like 1). A subset of neutrophils which clustered closest to monocyte/macrophages highly expressed the transcription factor *FOS* (c-Fos) (C). To investigate which immune cells were increased in the CNS in TBM in association with higher neutrophil counts, broad cell type proportions were compared between patients with low (≤5 cells/mm^3^) and high (>5 cells/mm^3^) CSF neutrophil counts from laboratory testing at the time of the lumbar puncture (LP). Patients with low laboratory neutrophil counts also had low numbers of cells annotated as neutrophils in the scRNAseq data. Patients with increased CSF neutrophils had significantly higher proportions of γδ/NK cells, proliferating cells, and Tregs, and lower proportions of B, plasma, cDC2 and MAIT cells (**Fig. S7E**).

**Fig. 4.**
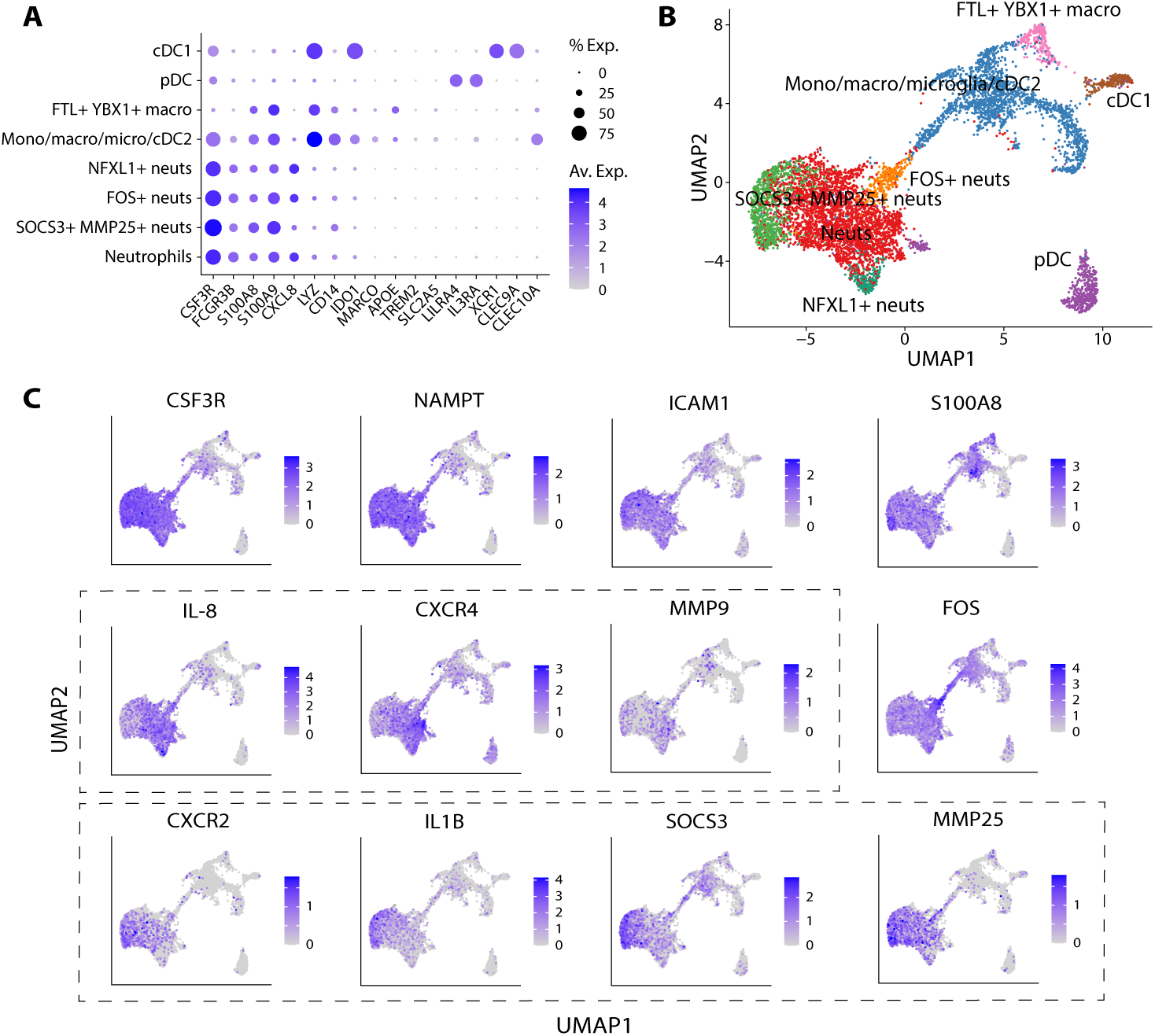
Neutrophils are highly activated and separate into IL-8 high and CXCR2 high phenotypes. **(A)** Dotplot of myeloid cell canonical gene expression of annotated clusters. **(B)** Annotated UMAP of 7,182 myeloid cells. **(C)** Feature plots of 12 myeloid marker genes with key differentiating genes from the two main phenotypes enclosed in dotted boxes. *CXCR2*, C-X-C motif chemokine receptor 2 or IL-8 receptor beta. IL-8, interleukin 8 (*CXCL8*); *CSF3R*, colony-stimulating factor 3 receptor; *NAMPT*, nicotinamide phosphoribosyltransferase (visfatin); *LYZ*, lysozyme; *APOE*, apolipoprotein E; *TREM2*, triggering receptor expressed on myeloid cells 2; *SLC2A5*, solute carrier family 2 member 5 (GLUT5); *CLEC10A*, C-type lectin domain family 10 member A; *CLEC9A*, C-type lectin domain family 9 member A; *XCR1*, XC motif chemokine receptor 1; *LILRA4*, leukocyte immunoglobulin-like receptor subfamily A member 4; *FOS*, c-Fos; *SOCS3*, suppressor of cytokine signaling 3; *MMP*, matrix metalloprotease; *NFXL1*, nuclear transcription factor, x-box binding like 1.

### Type 1 interferon associated genes were overexpressed in several cell types in microbiologically positive TBM

**Figure 3B** shows type 1 interferon-stimulated genes gene expression across all cells. Type 1 interferon-stimulated genes (ISG) were moderately expressed in the majority of CD8 and CD8 T cell subclusters, but were particularly highly expressed in two CD8 effector memory-like T cell clusters (ISG+ Tem/effector, IFI27+ Tem/effector) (**Fig. 2C**) and one CD4 T cell cluster (ISG+) (**Fig. 2F)**. As with cytotoxic genes, several cell types showed increased expression of genes associated with type 1 interferon signaling in patients with microbiologically positive TBM. ISG (*IFI27*, *IFIT3*, *IFI44L*, *ISG15*, *MX1*) were overexpressed in several clusters of CD8 cells in patients with microbiologically positive TBM, along with signal transducer and activator of transcription protein 1 (*STAT1*) (**Fig. 3D**). FCER1G+ NK cells and naïve-like γδ cells overexpressed the interferon regulatory factor *IRF8*, and the *IFI6* and *OAS3*, respectively, in microbiologically positive disease (D). In B cells, *STAT1*, *IFI6* and *IFI27* were overexpressed in IGHA^hi^ memory cells and IGKC+ plasma cells (D). In contrast, the EPHB6+ and KLRB1+ CD4 T cell subclusters overexpressed *GBP1* and *GBP2*, which initiate non-canonical caspase 4-mediated inflammasome activation and are associated with IFN-γ (*15*, *16*) (D). Overall, this suggests that type 1 interferon signaling is increased in several immune cell types in the CSF in patients with higher bacterial loads.

### Type 1 interferon associated gene expression increased despite antibiotic therapy and remained above pre-treatment levels for several weeks in the blood and CSF

To investigate interferon-associated gene expression changes over time in the CSF and blood in HIV-associated TBM, we performed longitudinal bulk RNA sequencing on lumbar CSF samples at two timepoints (≤day 7 and week 4) and blood samples at four timepoints (day 0, day 7, week 2 and week 8) taken from PLWHA with TBM in a separate cohort (**Table S7 to S10**). Laboratory counts of CSF lymphocytes, neutrophils, protein and glucose did not change significantly between participants between the first week of treatment and week 4, but blood neutrophil counts fell significantly over time (**Fig. S8**).

In the CSF, DEG between the first week post-enrolment and week 4 found the Th17 transcription factor *RORC*, the microglial marker *TREM2* and the glutamate transport excitatory amino acid transporter 1 (EAAT1, *SLC1A3*) overexpressed early in treatment, whilst *GNLY*, IL-8 receptor beta (*CXCR2*), IL-1 receptor antagonist (*IL1RN*), and matrix metalloprotease 9 (*MMP9*) were significantly overexpressed at week 4 (**Fig. 5A**). The interferon alpha response was the top upregulated pathway at week 4 in gene set enrichment analysis (GSEA) (**B**). Several type 1 ISG (*ISG15*, *MX1*, *IFIT1*, *IFIT3*, *IFI16*, *IFI35*) significantly increased by week 4 compared to the first week of study enrolment, along with *IRF7*, and IFN-γ-induced *GBP1* (**C**). In the blood, pathways associated with complement activation, phagocytosis and B cell receptor signaling were higher at baseline, whereas type 1 interferon signaling pathways were significantly repressed at day 0 compared with week 8 using GSEA (**D**). Expression of the same canonical ISG differentially expressed in the CSF peaked at either day 7 or week 2 depending on the gene (**E**). Overall, this suggests that type 1 interferon signaling increases on antibiotic therapy and is sustained above pre-treatment levels for at least one month in both the lumbar CSF and peripheral blood in HIV-associated TBM.

**Fig. 5.**
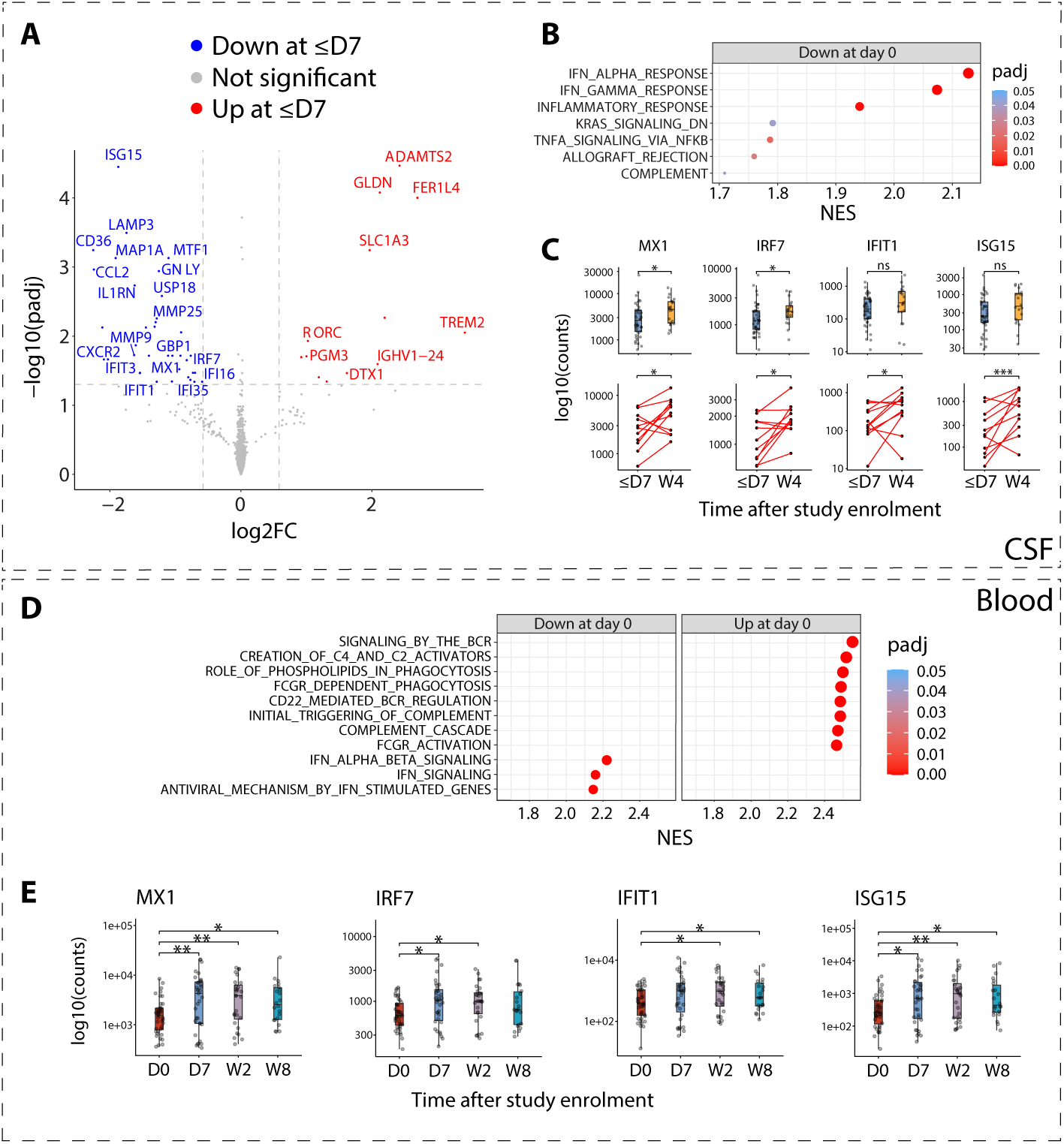
Gene expression associated with type 1 interferon is delayed and sustained in the lumbar CSF and blood in HIV-associated TBM. **(A)** Volcano plot of differentially expressed genes in the CSF between day 7 and week 4 post-enrolment. Red, overexpressed at day 7; blue, overexpressed at week 4. **(B)** GSEA showing significantly downregulated Hallmark pathways at day 7 compared to week 4 in the CSF. **(C)** Gene expression of four canonical type 1 interferon-associated genes at day 7 and week 4 in the CSF. The top row shows all samples with adjusted p values from unpaired linear modelling. The bottom row shows paired samples only with adjusted p values from paired linear modelling. **(D)** GSEA showing significantly down and upregulated Reactome pathways at day 7 compared to week 4 in the blood. **(E)** Gene expression of four canonical type 1 interferon-associated genes at enrolment, day 7, week 2 and week 8 in the blood.

## DISCUSSION

PLWHA have the highest chance of developing TBM, the highest risk of death and disability from TBM, and the fewest evidence-based options to moderate the immune response to ameliorate these poor outcomes. Our understanding of the immune dysregulation in TBM, particularly in PLWHA, is limited, impacting the ability to develop or repurpose novel host-directed therapies. This study characterised the CNS cellular immune response to HIV-associated TBM at a single cell level and identified several correlates of bacterial load that may deserve future evaluation either as diagnostics or as a means to track the efficacy of new treatments.

Combining all 25 patients into a single UMAP masks heterogeneity in CSF cell composition in HIV-associated TBM. Notwithstanding, CD8 T cells were always the predominant cell type, even in patients with normal CD4 counts. In certain patients, CD4 T cells, γδ/NK or B cells exceeded 20% of total cells. This heterogeneity must be considered when making inferences from low numbers of patients and may have implications for the development and stratification of host-directed therapies if the impact of immunomodulation depends on CSF cell composition. CD8 T cell predominance contrasts with previous scRNAseq studies in TBM and other inflammatory CNS conditions in which CD4 T cells predominate (*12*, *13*, *17*), and may be explained by depletion of CCR5+ and CXCR4+ CD4 T cells in PLWHA. CD4 T cells were either shifted towards a naïve-like phenotype with high *TCF7*, *IL7R*, *SELL* and *LEF1* expression, or were regulatory in phenotype. This naïve-like phenotype is expanded in PBMC of *M. tuberculosis*-experienced PLWHA compared to before HIV-1 infection (*18*), and is increased in the pericardial fluid of PLWHA with TB pericarditis compared to HIV-uninfected patients (*19*).

The lumbar CSF milieu was highly cytotoxic. The majority of CD8 T cells expressed a selection of cytotoxic genes, including granzymes and perforin. In humans, granzymes canonically help induce cell death of infected cells. *GZMA* and *GZMB* have been relatively well-characterised, but the function of other granzymes remains poorly defined, including their non-cytotoxic extracellular functions (*20*). In our data, *GZMA* and *GZMM* were widely expressed. *GZMB* expression was only seen in GNLY+ and IFNG+ clusters, whereas *GZMK* was widely expressed outside NK-like and GNLY+ clusters. CD8 T cells expressing high levels of *GZMK* are the main subset of CD8 T cells in inflamed tissues in several diseases (*21*). Unstimulated CD8 T cells constitutively express *GZMK*, fitting with this broad expression in our CD8 T cells, most of which may not be antigen specific. In contrast, GZMB is not released constitutively, but only in response to TCR stimulation (*22*). Tram *et al*. found an increased proportion of GZMK+ CD8 T cells in CSF compared with blood in their TBM scRNAseq data (*13*). GZMK was recently found to cleave the C2 and C4 proteins, with the resulting convertases activating complement, including the anaphylatoxins C3a and C5a, opsonins C4b and C3b, and the membrane attack complex (*22*). In our prior transcriptomic analysis, complement activation was associated with progressive HIV-TB with transcripts associated with the classical complement pathway overabundant in the blood in subclinical and active TB (*23*). In addition, GZMK stimulates LPS-CD14 binding and TLR signaling to potentiate the monocytic cytokine response; whether the highly lipid rich cell wall of *M. tuberculosis* induces this in CNS macrophages/microglia is worthy of further work (*24*). This cytotoxic environment was increased in patients with higher bacterial loads, driven by cytotoxic gene expression from CD4, MAIT and CD56^hi^ NK cells rather than CD8 T cells. Day *et al*. showed that polyfunctionality (defined by IFN-γ, IL-2 and TNF detection) of *M. tuberculosis*-specific CD4 T cells reduces in smear-positive PTB compared with smear-negative PTB and LTBI, and recovers with treatment (*25*, *26*). Instead, in smear-positive PTB, single-positive TNF-producing *M. tuberculosis*-specific CD4 T cells are increased. Together, this may mean that in the presence of higher bacterial loads, cytokine polyfunctionality is replaced by cytotoxicity in CD4 T cells in the CSF in TBM. Overall, the majority of CD8 T cells, the predominant cell type in HIV-associated TBM, were highly cytotoxic with low cytokine expression. These cells may not contribute directly to *M. tuberculosis* control, and could contribute to immunopathology through a positive feedback cycle of cytotoxicity and complement activation. *GZMK*, newly associated with complement activation, was widely expressed in CD8 T cells and may contribute to immunopathology in TBM. PLWHA may be more affected than HIV-uninfected patients because of an increased proportion of CD8 T cells and cytotoxicity mediated by CD4, NK and MAIT cells may be exacerbated further in patients with higher bacterial loads.

To our knowledge, this is the first study to identify and characterise neutrophils in TBM at a single cell resolution. Neutrophils are critical to understanding the immunopathology in TBM, associating with poor outcome in both CSF and blood (*10*). CSF neutrophils correlate with CSF *M. tuberculosis* bacterial load (*7*). In pulmonary TB, they provide a replicative niche for *M. tuberculosis* and cause tissue destruction. High pulmonary neutrophil counts in mice cause a necrotic granulomatous phenotype, which can be induced in mice with pre-existing *M. tuberculosis* immunity by depleting CD4 T cells, mimicking HIV infection. Neutrophil depletion using ⍺Ly6G or a CXCR2 inhibitor reduces lung bacterial load and cavitation (41). In our cohort, neutrophils did not differ in proportion between patients with and without microbiologically positive TBM but were highly activated. *NAMPT* and *ICAM1* were both highly expressed by, and specific to, neutrophils in our data. *NAMPT*, also known as visfatin, is expressed in response to cytokines and LPS, and inhibits neutrophil apoptosis (*27*). *ICAM1* expression in neutrophils is increased by LPS, TNF, GM-CSF, and lymphotoxin-alpha (LT⍺) *in vitro*, and neutrophils with increased *ICAM1* expression have enhanced phagocytic capacity (*28*). In mice, neutrophil *ICAM1* expression correlates with increased phagocytosis and reactive oxygen species generation, and ICAM1 deficiency causes defective phagocytosis (*29*). Phenotypically, neutrophils split broadly into MMP25+ IL1B+ SOCS3+, and CXCL8+ clusters. SOCS3 is a negative regulator of G-CSF signaling and neutrophil-specific Socs3-deficient mice develop autoimmune encephalitis (*30*, *31*). *SOCS3* expression in combination with *MMP25* and *IL1B* may suggest an activated phenotype which has begun to induce *SOCS3* to contain inflammation. The other cluster expressed CXCL8 (IL-8), a neutrophil chemoattractant, suggesting that these neutrophils may drive further neutrophil recruitment. Together, this suggests that neutrophils in the CSF in TBM may be inhibited from apoptosis, drive further peripheral neutrophil recruitment, and have increased effector functions. Inhibiting CNS neutrophil chemotaxis and ingress, for example by CXCR2 inhibition, may be a potential strategy to abrogate this ‘feed forward’ loop and reduce intracranial inflammation.

Type 1 IFN is associated with impaired control of *M. tuberculosis* infection, and excess signaling in the CSF may contribute to immunopathology in TBM. Type 1 IFN suppresses host-protective cytokines, impairs the Th1 response, and limits prostaglandin E2 (PGE2), which promotes apoptosis rather than necrosis of macrophages (*32*). In addition, type 1 IFN represses responsiveness to the antibacterial effects of IFN-γ (*33*). Several Ifnar1^-/-^ mouse studies have shown reduced *M. tuberculosis* bacterial loads in the lung and/or improved survival (*32*, *34–36*). Induction of type 1 IFN in mice impairs control of bacterial growth and exacerbates lung immunopathology (*37–39*). In our data, there was a cluster of CD8 effector memory-like cells that highly expressed type 1 interferon-stimulated genes (ISG). This ISG+ cluster was also seen in CD4 T cells and did not otherwise appear phenotypically different to neighbouring non-ISG+ cells. Patients with microbiologically positive disease had increased expression of ISG across most cell types, including CD8 T, γδ, NK, MAIT, dendritic and B cells. The delayed peak on treatment in type 1 interferon response shown in our bulk transcriptomic data contrasts with the response seen in pulmonary TB. Amongst TB contacts who progressed to active pulmonary TB, a gene module in the blood associated with interferon signaling peaked pre-treatment and resolved with antibiotic therapy (*40*). In the blood in our HIV-associated TBM cohort, type 1 interferon signaling gene transcripts were still increased at week 8 compared to baseline. In the CSF, the same pattern was seen, with an increase from the first week to week 4. This sustained response may explain why CNS inflammation persists for so long despite treatment in TBM and why CSF neutrophils, which are recruited in response to type 1 interferon, remain high at week 4. However, the reverse may occur; neutrophils are an important source of type 1 interferon in TB (*41*), and it may be persistently raised neutrophils that are driving sustained type 1 interferon secretion. Overall, type 1 IFN is detrimental in the immune response to TB, and in HIV-associated TBM there are distinct cell clusters responding to type 1 IFN signaling, which associate with bacterial load.

There are limitations to this study. TBM patients with low cell counts in their CSF could not be included because of the minimum number of cells required for the 10x Flex. This risks biasing results towards patients with higher cell counts who may not be representative of TBM patients as a whole. The 10x Flex is probe-based, which means that only information on the ∼18,000 human genes targeted by the probes is available. Variable genes, such as human leukocyte antigen (HLA) genes, which can help identify *M. tuberculosis*-specific T cells, and *TRAV1-2* genes which can identify MAIT cells, were not available. Probe-based sequencing meant that TCR and BCR sequencing was also not available, so inferences about clonality could not be made. Our cohort had relatively mild disease since unwell patients often die before day 7 or LP is contraindicated. Mild disease severity and ongoing study blinding meant that comparisons based on outcome were not possible. All patients received a weaning corticosteroid regimen as per international guidelines. Corticosteroids influence the immune response, and this influence may have differed between patients and impacted gene expression in the longitudinal bulk RNA sequencing. Some heterogeneity in bacterial load and host immune response may be explained by the duration of disease prior to presentation which cannot be reliably measured or controlled for. Patients received a median of 10 days’ antibiotic treatment at the time of LP. However, immune resolution is slow in TBM, and it is unlikely that dramatic changes occur during this early treatment period. This study did not include healthy or non-TBM controls. Enrolling healthy controls is challenging in this context because an LP is an invasive procedure and therefore comes with ethical concerns and healthy CSF is paucicellular. Non-TBM controls such as cryptococcal meningitis or viral meningitis must be recruited within a study setting but are not treated with corticosteroids making comparisons difficult. Finally, 11/25 TBM patients had probable or possible TBM and may have had a diagnosis other than TBM. This may have biased the results of comparisons between those with and without microbiological confirmation. It is impossible to completely rule out an alternative infection or co-infection, but all patients recovered during specific TBM treatment, except for one possible TBM participant who died.

In conclusion, this study is the largest scRNAseq study to date in TBM, including 188,983 cells from the CSF of 25 adults with HIV-associated TBM. Our findings suggest a hyper-inflammatory immune response in the CSF in HIV-associated TBM characterised by an overabundance of granzyme rich cytotoxic bystander CD8 T cells, ‘feed-forward’ recruitment of highly activated neutrophils, and sustained suppression of protective immunity via excess type 1 IFN signaling, all of which is exacerbated by higher bacterial loads. Bacterial load, neutrophils and type 1 interferon inversely correlate with protective CD4 T and B cell adaptive immunity and this dysregulated response is likely to be more severe in PLWHA who have a reduced proportion of CD4 T cells. The relative contributions of bystander and *M. tuberculosis*-specific cells in mediating inflammatory pathology and the drivers of CNS neutrophil ingress in TBM require further study.

## MATERIALS AND METHODS

### Study approval

Collection and analysis of INTENSE-TBM samples was approved by the University of Cape Town (UCT) Human Research Ethics Committee as part of the INTENSE-TBM multiomic sub-study (004/2020). The LASER-TBM trial, including permission to collect and analyse samples, was approved by the UCT Human Research Ethics Committee (293/2018), Walter Sisulu University Human Research Committee (012/2019), and the South African Health Products Regulatory Authority (20180622). HIATUS-3 was approved by the University of Cape Town (UCT) Human Research Ethics Committee (207/2020). All participants provided written informed consent following explanation of the nature and possible consequences of the studies in their first language.

### CSF collection, processing and sequencing

CSF was obtained from an LP performed at day 7 following enrolment as part of the INTENSE-TBM trial (*42*). Up to 6 ml CSF was collected in a 15 ml falcon tube at the bedside after prioritising necessary clinical investigations. Samples were transferred to the biosafety level 3 laboratory at 4°C and processed within two hours of collection. Fresh CSF was centrifuged at 400 RCF for 10 minutes at 4°C and supernatant removed. Cells were fixed in a 4% formaldehyde fixative solution, as described in the Demonstrated Protocols CG000478 and CG000553 and using Chromium Next GEM Single Cell Fixed RNA Sample Preparation Kit (10x Genomics PN-1000414), then stored at -80°C for up to 12 months. Fixed samples were processed following the Chromium Fixed RNA Profiling Reagent Kits for Multiplexed Samples Protocol (10x Genomics CG000527, Revision A) using the Chromium Fixed RNA Kit, Human Transcriptome, 4 rxns x 4 BC (10x Genomics PN-1000475) and 4 rxns x 16 BC (PN-1000476). Gene expression was measured using barcoded probe pairs designed to hybridise to mRNA specifically. Using a microfluidic chip, the fixed and probe-hybridised single cell suspensions were partitioned into nanolitre-scale Gel Beads-in-emulsion (GEMs). A pool of ∼737,000 10x GEM Barcodes was sampled separately to index the contents of each partition. Inside the GEMs, probes were ligated and the 10x GEM Barcode was added, and all ligated probes within a GEM share a common 10x GEM Barcode. Barcoded and ligated probes were then pre-amplified in bulk, after which gene expression libraries were generated (User Guide: CG000527, Chromium Fixed RNA Profiling Reagent Kits for Multiplexed Samples) and sequenced (NovaSeq 6000, NovaSeq X. Sequencing read configuration: 28-10-10-90). Samples were processed and sequenced in three batches. All samples in the same batch were multiplexed using sample-specific probe sets and sequenced together. Two samples in batch 3 were double barcoded to maximise use of the 16-plex kit. These samples therefore had twice the number of cells sequenced.

### Ambient RNA and doublet removal

**Fig. S9** shows a flowchart of the bioinformatics analysis. CellRanger v9.0.1 filtered matrices, aligned to the human GRCh38 genome, were processed in R v4.4.1 using Seurat v5.2.1. Ambient RNA contamination and predicted doublets were removed using decontX v1.2.0 (*43*) and scDblFinder v1.18.0 (*44*), respectively, with default settings. The median predicted RNA contamination percentage per cell was 0.54% (range 0-99.98%) (**Table S5**). The median percentage of doublets predicted was 8.42% (range 5.63%-12.42%) (**Table S6**).

### Filtering out poor quality cells

Histograms of samples with known neutrophil populations showed a bimodal distribution, with neutrophils forming a peak with a lower number of genes/cell (**Fig. S10**). A conventional cut off would have excluded many of these neutrophils. Therefore, a cut off of 100 genes/cell was used to preserve neutrophils. Overall, the percentage of mitochondrial RNA per cell was low. In most samples, almost all cells had <2.5% mitochondrial RNA, but two samples had a significant proportion of cells between 2.5-5% so a cut off of >5% was used to exclude cells from downstream analysis (**Fig. S11**). Low quality cells with <100 genes and >5% mitochondrial reads were therefore excluded. Violin plots of the number of genes, unique molecular identifiers (UMI) and mitochondrial RNA percentage per cell for the 25 individual samples before and after filtering are shown in **Fig. S12.** The median number of genes per cell was higher for predicted doublets than singlets in all samples, but there was no difference in percentage of mitochondrial genes per cell. Scaling, normalisation and variance-stabilisation were performed using the SCTransform function using default settings. Principal component analysis (PCA), nearest neighbour calculations, uniform manifold approximation and projection (UMAP) and Louvain clustering were performed using the Seurat packages RunPCA(), FindNeighbors(), RunUMAP() and FindClusters() respectively using the top 2,000 variable features and first 30 principal components.

### Integration using reciprocal PCA

All samples were merged and integrated using reciprocal principal component analysis (RPCA) in Seurat’s IntegrateLayers() function with the default k.anchor = 5. RPCA projects each dataset into the others’ PCA space and identifies anchors using mutual nearest neighbours, then uses these anchors to integrate the data sets. Nearest neighbour, UMAP and clustering were repeated with the new integrated PCA embeddings using the top 30 dimensions and compared with the unintegrated UMAP to ensure batch differences had been removed (**Fig. S13**). Seurat’s FindSubCluster() was used to further subcluster one cluster of mixed cell types.

### Clustering and annotation

Louvain clustering and annotation were done on integrated UMAP embeddings. FindClusters() was run at a selection of resolutions to explore the optimal resolution. Resolution 0.9, which contained 37 clusters, was the lowest resolution which preserved myeloid cell types and was selected for subsequent subclustering and annotation (**Fig. S14**). Despite doublet removal, cluster 35 (230 cells) showed a mix of T and B cell expression from all clusters, suggesting it consisted of doublets. This cluster was removed leaving 36 clusters. Cluster annotation was done using canonical immune cell marker expression, the top differentially-expressed genes between clusters using the FindAllMarkers() Seurat function, and automatic cell labelling using the SingleR v2.8.0 (Monaco, Blueprint and Human Primary Cell Atlas references) and Celltypist v1.7.1 (Immune Low reference) packages (*45*, *46*). Canonical cell markers were taken from the literature. In the case where a cell (sub)type was unclear, differentially expressed genes for that particular cluster were used to inform the cell type or function of that cluster. The UMAP contained a spatially distinct group of cells which clustered with cluster 26 at resolution 0.9. Even at a resolution of 2.0, this cluster did not separate. These cells highly expressed the canonical MAIT cell markers ZBTB16 (zinc finger and BTB domain-containing protein 16 [promyelocytic leukaemia zinc finger, PLZF]), KLRB1 (killer cell lectin-like receptor subfamily B, member 1, CD161), and SLC4A10 (solute carrier family 4 member 10). TRAV1-2 was not included in the probe set used for the 10x Flex because it is a variable domain. Automated tools labelled the cells as mucosal-associated invariant T (MAIT) cells, with very high confidence. To separate these putative MAIT cells, the Seurat FindSubCluster() function was used at a resolution of 0.2 to subcluster cluster 26 into three subclusters. Cluster 26_0 shared expression with NK/γδ cells, 26_1 was the MAIT cell cluster and 26_2 mapped to the region identified as progenitors by all four automated annotation tools (**Fig. S15**).

Coarse cell types were subsetted and re-analysed to explore their phenotype in more detail. After subsetting NK and T cells by removing myeloid, neutrophil, B cell and plasma cell clusters, CD8 T cells were identified as cells expressing any CD3 gene (CD3D or CD3E or CD3G) *and* CD8A. CD4 T cells were identified as cells expressing any CD3 gene (CD3D or CD3E or CD3G) *and* CD4. NK and γδ cells, plasma and B cells and myeloid cells were subsetted from their respective clusters in **Fig. 1E**. Raw counts from these subsetted cell types were re-normalised, merged, integrated and clustered. Cluster annotation of subsetted cells was performed using a combination of canonical markers and differentially expressed genes as outlined above.

### Composition analysis and differential gene expression of scRNAseq data

Differential abundance of cell populations between clinical phenotypes was performed using sccomp v1.10.0, which uses a generalised method for differential composition and variability analyses based on sum-constrained independent β-binomial distributions that outperforms other methods (*47*). Estimation was done through Hamiltonian Monte Carlo via the Bayesian inference framework Stan (*48*). The analysis was corrected for age and sex. Pseudobulk DEG between those with and without microbiologically positive TBM was done using the Wald test in DESeq2 within the muscat package (*49*). Pseudobulk analysis was done by aggregating the data into per-group clusters. Pseudobulks with low numbers of cells were excluded. As with the composition analysis, age and sex were modelled as co-variates.

### Blood bulk RNA sequencing

Whole blood was collected directly into PAXgene RNA tubes (QIAGEN/BD Biosciences) from 40 patients enrolled into the LASER-TBM trial (*50*). Demographic data for the blood RNA sequencing are shown in Tables S7 and S8. Total RNA was extracted using the PAXgene Blood RNA Kit (QIAGEN, Venlo, Netherlands). Samples were stored at -80°C before and after RNA extraction. Quantity and quality of RNA was assessed using a Qubit fluorimeter (ThermoFisher) and Agilent 2100 Bioanalyzer. Samples with RNA integrity number (RIN) <7 were excluded. RNA libraries were constructed using Tecan’s Universal Human Blood RNA-Seq kit. Ribosomal and globin RNA were removed using Tecan’s AnyDeplete system. DNA library size distribution was assessed using the D1000 TapeStation kit processed on the 4200 TapeStation (Agilent Technologies). The data were analysed by Agilent’s 4200 TapeStation Analysis Software. Concentration was measured using the QuantiFluor dsDNA System (Promega Corporation). Libraries were sequenced on an Illumina HiSeq4000 with a single-end 100 cycle read configuration.

Raw data were processed using nf-core/rnaseq (v3.10.1) of the nf-core collection of workflows. This workflow used FASTQ files as input, performed quality control (QC), trimming and (pseudo-)alignment to the GRCh38 release 95 human reference genome, and produced a gene expression matrix and QC report (*51*). The pipeline was executed with Nextflow (v22.10.3) (*52*). The output matrix was read into R (v4.2.0) in which subsequent analysis was done (*53*).

Genes with raw counts ≥5 in ≥5 samples were used for differential gene expression analysis using DESeq2 (*54*). DESeq2 then uses the apeglm package to perform generalised linear modelling to estimate log fold changes (LFC) for each gene between groups. Raw LFC shrinks LFC with high variability and low counts towards zero to reduce noise (*55*). Paired timepoint analysis was done by including patient number in the model. Normalisation for principal component analysis and heatmaps was done using variance stabilising transformation (VST). *In silico* deconvolution was done using LM22, a validated gene signature matrix of 22 cell types using CIBERSORTx (*56*). Gene set enrichment analysis (GSEA) was done using genes ranked by log2FC output from DESeq in the fgsea package (v.1.29.1) using the Reactome gene set.

### CSF bulk RNA sequencing

CSF was collected directly into PAXgene RNA tubes (QIAGEN/BD Biosciences) at the bedside from patients enrolled in the LASER-TBM and INTENSE-TBM trials (*42*, *50*). A single patient with definite TBM was included from HIATUS-3, an observational study. Demographic data for the CSF RNA sequencing are shown in Tables S9 to S10. Lumbar puncture was done at day 0 (HIATUS-3), day 3 (LASER-TBM) and day 7 (INTENSE-TBM), and week 4 (LASER-TBM and INTENSE-TBM) per the study protocols. Total RNA was extracted using the PAXgene Blood RNA Kit (QIAGEN, Venlo, Netherlands). RNA extraction of samples from the two trials was done at separate timepoints but DNA libraries were prepared at the same time. Samples were stored at -80°C before and after RNA extraction. The quantity and quality of RNA was assessed using a Qubit fluorimeter (ThermoFisher) and Agilent 2100 Bioanalyzer. Concentrations of total RNA were low (median 1.12 ng [range 0.494-403 ng]). The Watchmaker DNA library prep kit (Watchmaker Genomics, Boulder, USA, CAT# 7K0102) with ribosomal depletion was used to construct libraries. CSF samples were divided into three batches of different input concentrations (2 ng [n = 36], 1 ng [n = 27], 0.5 ng [n = 46]) so that RNA loading could be maximised for samples with higher RNA concentrations. Two samples were excluded from the library prep because they had concentrations < 0.5 ng. Following library preparation, DNA library size distribution was assessed using the D1000 TapeStation kit processed on the 4200 TapeStation (Agilent Technologies). The data were analysed by Agilent’s 4200 TapeStation Analysis Software. Concentration was measured using the QuantiFluor dsDNA System (Promega Corporation). Libraries were sequenced on an Illumina NovaSeq 6000 with a paired-end 100 cycle read configuration with a target sequencing depth of 30 million reads/sample. After the first round of sequencing, 52/120 samples had enough reads, 38/120 were re-pooled for top up sequencing, and 30/120 were excluded because of a lack of reads. In total, five rounds of re-pooling and sequencing were done, which achieved a median read depth of 41 million reads/sample (range 16.2-149 million) in the 82 included samples.

Raw data were processed as per the blood RNA sequencing. Genes with raw counts ≥5 in ≥5 samples were used for differential gene expression analysis using DESeq2. Paired timepoint analysis was done by including patient number in the model. All modelling included ‘batch’ to account for differences between the different library prep RNA input concentrations. Functional enrichment analysis was done using the same methods as the blood transcriptomics.

### Statistics

Parametric continuous data were compared using t-test if two groups, or ANOVA if more than two groups. Non-parametric continuous data were compared using unpaired Mann Whitney U if two groups, or Kruskal-Wallis rank sum test if more than two groups. Categorical data were compared using the Chi-squared test.

## Acknowledgments

We would like to acknowledge all study participants and staff, especially Rene Goliath, Mpumi Maxebengula, Sheena Ruzive, Laeeqa Allie, Lauren Barron, and Petro Booysen. We are grateful to the Western Cape province for the use of its facilities and consultants, in particular Trevor Mnguni, Tom Crede, Patrick Szymanski, Yacoob Vallie, Ismail Banderker, and Muhammed S Moosa. In addition, we thank the INTENSE-TBM consortium, and Asma Toefy for supervision of Nonzwakazi Bangani. Finally, we would like to thank the Genomics and Bioinformatics and Biostatistics (BABS) Science and Technology Platforms at the Francis Crick Institute for their technical support, in particular Maria Rodriguez-Lopez and Gavin Kelly.

## Author contributions

Conceptualization: JRB, RJW

Data curation: JRB, AGD

Formal analysis: JRB, CB

Methodology: JRB, RJW

Investigation: JRB, NB, HS, CJS, GM, FB, AGD

Visualization: JRB

Funding acquisition: RJW, FB, RL

Project administration: CJS

Supervision: RJW

Writing – original draft: JRB

Writing – review & editing: RJW, AOG, DLB, AGD, MG, BR, RL, KAW, JRB

## Data and materials availability

Raw fastq files and processed CSF scRNAseq data, CSF bulk RNAseq and blood bulk RNAseq are deposited in the GEO at accessions GSE321718, GSE324044 and GSE324265, respectively. Transfer of CSF and blood for further analysis is subject to ethical approval and material transfer agreements.

## Supplementary figures

**Fig. S1. All cells. (A)** Unannotated UMAP of the 188,983 cells after predicted doublet and contamination removal at resolution 0.9. **(B)** Dot plot of original clusters classed as B or plasma cells based on canonical markers. **(C)** Dot plot of original clusters classed as myeloid cells based on canonical markers. **(D)** Dot plot of original clusters classed as NK or T cells based on canonical markers. **(E)** Pie chart of proportions of coarse cell types after manual and automated annotation. **(F)** Comparison of cell composition by microbiological-confirmation using the sccomp R package adjusting for age and sex. The black boxplot is the observed data; the blue boxplot is simulated data using a posterior predictive distribution. **(G)** Comparison of cell composition by ART at enrolment using the sccomp R package adjusting for age and sex.

**Fig. S2. Correlation matrix of cell types**. Correlation matrix of absolute cell counts across the 25 patients calculated with Spearman’s rank correlation. Laboratory plasma CD4 count (cells/mm^3^) and HIV-1 viral load (copies/ml) highlighted in bold. Blue represents positive correlation and red negative correlation. Cell types ordered by hierarchical clustering.

**Fig. S3. CD8 T cells**. **(A)** Dot plot of top 5 DEG per cluster. **(B)** Dot plot of canonical genes used to guide manual annotation. **(C)** Unannotated clustering of CD8+ T cells. Raw counts from 123,557 CD8A+ CD3+ cells were re-normalised, merged, integrated and clustered at resolution 0.8 to create 27 clusters. **(D)** Pie chart of CD8 T cell cluster proportions. **(E)** Comparison of CD8 T cell subcluster composition by microbiological-confirmation using the sccomp R package adjusting for age and sex. The black boxplot is the observed data; the blue boxplot is simulated data using a posterior predictive distribution. GXPU, GeneXpert Ultra. *IFNG*, interferon gamma; *GNLY*, granulysin; Tem, effector memory T cell.

**Fig. S4. CD4 T cells**. **(A)** Dot plot of top 5 DEG per cluster. **(B)** Dot plot of canonical genes used to guide manual annotation. **(C)** Unannotated UMAP of CD4 T cells. Raw counts from 21,844 CD4+ CD3+ cells were re-normalised, merged, integrated and clustered at resolution 0.4 to create 12 clusters. **(D)** Pie chart of CD4 T cell cluster proportions. **(E)** Comparison of CD4 T cell subcluster composition by microbiological-confirmation using the sccomp R package adjusting for age and sex. The black boxplot is the observed data; the blue boxplot is simulated data using a posterior predictive distribution. **(F)** Volcano plot of DEG of CXCR6+ CCL5+ GZMA+ IFNG+ CD4 T cell by microbiological confirmation. **(G)** Gene set enrichment analysis of DEG by microbiological confirmation in the CXCR6+ CCL5+ GZMA+ IFNG+ CD4 T cell cluster done using the Gene Ontology Biological Processes gene set, ranked by log2 fold change. ISG, interferon-stimulated gene; *KLRB1*, killer cell lectin like receptor 1 (CD161); *EPHB6*, ephrin type-B receptor 6; *GZMA*, granzyme A; *IFNG*, interferon gamma.

**Fig. S5. NK and γδ cells. (A)** Dot plot of top 5 DEG per cluster. **(B)** Dot plot of canonical genes used to guide manual annotation. **(C)** Unannotated UMAP of NK and γδ T cells. Raw counts from 11,483 NK and γδ cells were re-normalised, merged, integrated and clustered at resolution 0.2 to create seven clusters. **(D)** Comparison of NK and γδ T cell subcluster composition by microbiological-confirmation using the sccomp R package adjusting for age and sex. The black boxplot is the observed data; the blue boxplot is simulated data using a posterior predictive distribution. **(E)** Volcano plot of DEG for the FCER1G+ NCAM1+ NK cell cluster by microbiological confirmation. *FCER1G*, Fc fragment of IgE receptor, gamma chain; *NCAM1*, neural cell adhesion molecule 1.

**Fig. S6. B and plasma cell subclustering by microbiological confirmation. (A)** Dot plot of top 5 DEG per cluster. **(B)** Dot plot of canonical genes used to guide manual annotation. **(C)** Unannotated UMAP of B and plasma cells. Raw counts from 14,042 B cells and plasma cells were re-normalised, merged, integrated and clustered at resolution 0.3 to create 11 clusters. **(D)** Annotated UMAP of 14,042 plasma and B cells. **(E)** Table of absolute numbers and proportions of B and plasma cell clusters. **(F)** Feature plots of immunoglobulin gene expression. **(G)** Comparison of B and plasma cell subcluster composition by microbiological-confirmation using the sccomp R package adjusting for age and sex. The black boxplot is the observed data; the blue boxplot is simulated data using a posterior predictive distribution. *IGHD*, IgD heavy chain; *IGHM*, IgM heavy chain; *IGHG1*, IgG heavy chain 1; *IGHA1*, IgA heavy chain 1; *IGKC*, immunoglobulin kappa light chain constant; *IGLC1*, immunoglobin lambda light chain constant 1.

**Fig. S7. Myeloid cells. (A)** Dot plot of top 5 DEG per cluster. **(B)** Dot plot of canonical genes used to guide manual annotation. **(C)** Unannotated UMAP of myeloid cells. Raw counts from 7,182 myeloid cells were re-normalised, merged, integrated and clustered at resolution 0.2 to create nine clusters. **(D)** Feature plots of myeloid cells with neutrophil canonical markers. **(E)** Comparison of coarse cell composition by laboratory CSF neutrophil count (high = ≥5 cells/mm^3^, low = <5 cells/mm^3^) using the sccomp R package adjusting for age and sex. The black boxplot is the observed data; the blue boxplot is simulated data using a posterior predictive distribution. *CSF3R*, colony-stimulating factor 3 receptor; *NAMPT*, nicotinamide phosphoribosyltransferase (visfatin); *LYZ*, lysozyme; *APOE*, apolipoprotein E; *TREM2*, triggering receptor expressed on myeloid cells 2; *SLC2A5*, solute carrier family 2 member 5 (GLUT5); *CLEC10A*, C-type lectin domain family 10 member A; *CLEC9A*, C-type lectin domain family 9 member A; *XCR1*, XC motif chemokine receptor 1; *LILRA4*, leukocyte immunoglobulin-like receptor subfamily A member 4.

**Fig. S8. Cell counts over time for participants in CSF and blood bulk RNA sequencing.** Comparison between early (≤7 days, n = 44) and late (week 4, n = 20) laboratory lymphocyte counts **(A)**, neutrophil counts **(B)**, protein **(C)** and glucose **(D)** for participants enrolled in INTENSE-TBM and LASER-TBM whose samples were used for CSF bulk RNA sequencing (patients = 53, samples = 64). +1 added to CSF cell counts, and +0.1 added to CSF protein to allow for log2 transformation of zero values. Comparisons between laboratory white cell count **(E)** and neutrophil count **(F)** at day 0 (n = 40), day 7 (n = 35), week 2 (n = 28) and week 8 (n = 22) for participants enrolled in LASER-TBM whose samples were used for blood bulk RNA sequencing (patients = 42, samples = 125). Comparisons made using Mann Whitney U to compare the two CSF timepoints and ANOVA to compare the four blood timepoints. ns = non-significant.

**Fig. S9. 10x Flex single cell RNA sequencing bioinformatics workflow.** Doublets refer to two or more cells that have been captured in the same droplet at the cell separation step, and therefore share the same cellular barcode. SCTransform refers to a normalisation and variance stabilisation technique using regularised negative binomial regression. RPCA, reciprocal principal component analysis.

**Fig. S10. Histograms of genes per cell**. Histogram of each sample plotting the number of genes per cell (nFeature_RNA) vs. frequency. Red dotted line indicates the cut off of 100 genes/cell used for filtering. Each panel is one donor.

**Fig. S11. Histograms of percentage of mitochondrial RNA per cell**. Histogram of each sample showing the percentage of mitochondrial RNA per cell vs. frequency. Red dotted line indicates the cut off of 5% mitochondrial RNA used for filtering. Each panel is one donor.

**Fig. S12. Genes, UMI and percent mitochondrial genes per cell before and after filtering.** A) Genes, UMI and percent mitochondrial genes per cell before filtering B) Genes, UMI and percent mitochondrial genes per cell after filtering cells with <100 genes and >5% mitochondrial RNA.

**Fig. S13. Reciprocal PCA integration. (A)** Unintegrated UMAP coloured by sample. **(B)** Unintegrated UMAP coloured by default clustering. **(C)** Reciprocal PCA integrated UMAP coloured by sample. **(D)** Reciprocal PCA integrated UMAP coloured by clustering at resolution 0.9. N.B. Sample colours are added by layer, with NS071 added last, meaning that later samples appear more prominent.

**Fig. S14. Integrated UMAPs at selected cluster resolutions**. UMAP at six resolutions using FindClusters(). Resolution 0.9 was used for annotation.

**Fig. S15. Annotation of MAIT cells. (A)** UMAP highlighting *ZBTB16* (zinc finger and BTB domain-containing protein 16, promyelocytic leukaemia zinc finger, PLZF) gene expression. **(B)** UMAP highlighting *KLRB1* (killer cell lectin-like receptor subfamily B, member 1, CD161). **(C)** Celltypist confidence scores for cells labelled as MAIT cells in data. Red means high confidence, blue means low confidence. **(D)** UMAP of all cells after subclustering cluster 26 using FindSubClusters() into three smaller clusters. Cluster 26_1 (n = 501) was re-labelled as MAIT cells based on the Celltypist confidence and canonical gene expression.

## Supplementary tables

**Table S1. Patient demographics for CSF single cell RNA sequencing split by microbiological status.** Continuous data are shown as median (IQR), categorical as number (%). Data compared between TBM patients with and without microbiologically positive disease (GXPU or *M. tuberculosis* culture positive) using unpaired Mann Whitney U for continuous data and Chi-squared for categorical data. Viral load (VL) measured in copies/ml, CD4 count measured in cells/µl. GXPU, GeneXpert Ultra; BMRC, British Medical Research Council; ART, antiretroviral therapy; CSF, cerebrospinal fluid; IQR, inter-quartile range.

**Table S2. Individual clinical data for the 25 patients in the scRNAseq study**. GXPU, GeneXpert Ultra; HIV, human immunodeficiency virus; CSF cerebrospinal fluid. IRIS, immune reconstitution inflammatory syndrome. CSF data are from the hospital laboratory reports from the same lumbar puncture as the scRNAseq cells were taken.

**Table S3. Cells removed during quality control**. UMI, unique molecular identifier; QC, quality control.

**Table S4. Cell type proportions per patient**. Absolute number and proportion of main cell types per patient compared with CD4 count, ART at enrolment and microbiological status.

**Table S5. RNA contamination per sample calculated by decontX.** Contamination estimated using decontX with default settings. Mean and median cell contamination per sample and across all samples shown. SD, standard deviation.

**Table S6. Predicted doublets per sample by scDblFinder**. Doublets predicted using scDblFinder with an estimated doublet rate of 7.5% and otherwise default settings. Absolute number of predicted cells and percentage of total cells predicted to be doublets shown, with mean and median for all samples. SD, standard deviation.

**Table S7. Patient demographics of blood bulk RNA sequencing cohort**. Continuous data are shown as median (IQR), categorical as number (%). Viral load (VL) measured in copies/ml, CD4 count measured in cells/µl. GXPU, GeneXpert Ultra; BMRC, British Medical Research Council; ART, antiretroviral therapy; CSF, cerebrospinal fluid; IQR, inter-quartile range.

**Table S8. Individual sample data from blood bulk RNA sequencing cohort**. Viral load (VL) measured in copies/ml, CD4 count measured in cells/µl. soc, standard of care.

**Table S9. Patient demographics of CSF bulk RNAseq sequencing cohort**. Continuous data are shown as median (IQR), categorical as number (%). Viral load (VL) measured in copies/ml, CD4 count measured in cells/µl. GXPU, GeneXpert Ultra; BMRC, British Medical Research Council; ART, antiretroviral therapy; CSF, cerebrospinal fluid; IQR, inter-quartile range.

**Table S10. Individual sample data from CSF bulk RNA sequencing cohort**. Viral load (VL) measured in copies/ml, CD4 count measured in cells/µl. soc, standard of care.

