## Supplemental Figures for "Contribution of cytotoxic CD8 T cells, neutrophils and type 1 interferon signaling to hyperinflammation in HIV-associated TB meningitis"

**Supplementary figures:**

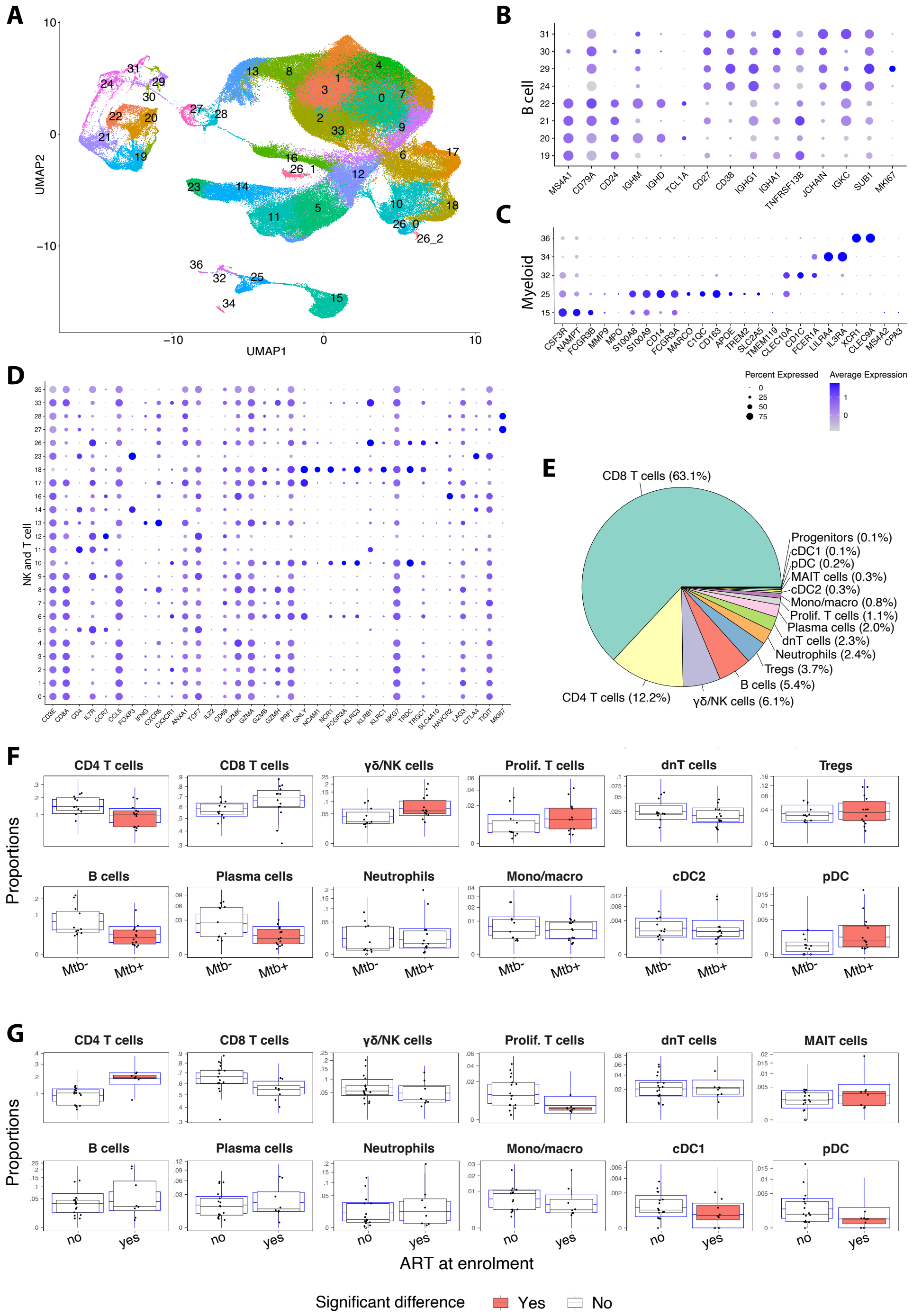


**Fig. S1. All cells. (A)** Unannotated UMAP of the 188,983 cells after predicted doublet and contamination removal at resolution 0.9. **(B)** Dot plot of original clusters classed as B or plasma cells based on canonical markers. **(C)** Dot plot of original clusters classed as myeloid cells based on canonical markers. **(D)** Dot plot of original clusters classed as NK or T cells based on canonical markers. **(E)** Pie chart of proportions of coarse cell types after manual and automated annotation. **(F)** Comparison of cell composition by microbiological-confirmation using the sccomp R package adjusting for age and sex. The black boxplot is the observed data; the blue boxplot is simulated data using a posterior predictive distribution. **(G)** Comparison of cell composition by ART at enrolment using the sccomp R package adjusting for age and sex.

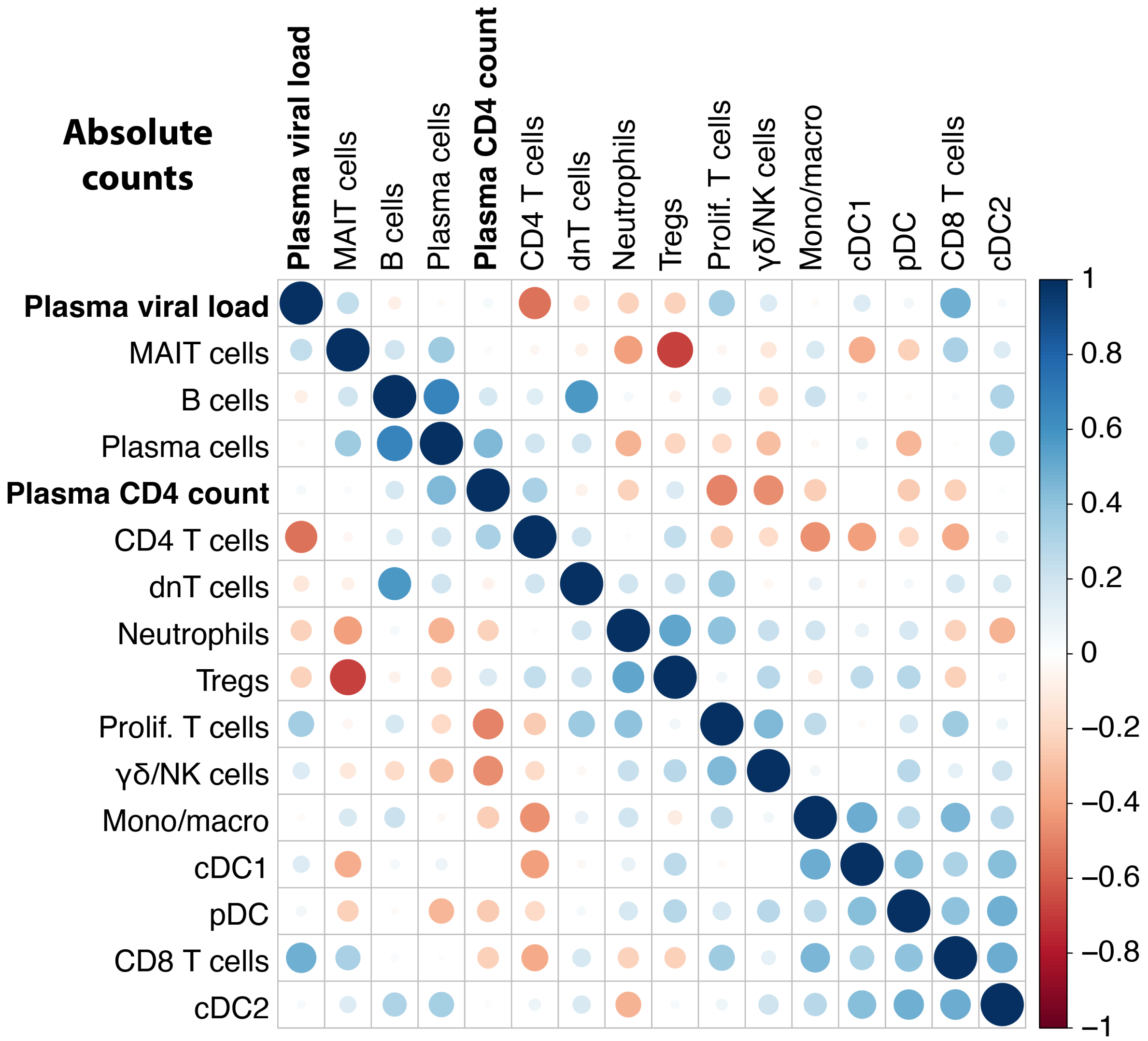


**Fig. S2. Correlation matrix of cell types**. Correlation matrix of absolute cell counts across the 25 patients calculated with Spearman's rank correlation. Laboratory plasma CD4 count (cells/mm$^3$) and HIV-1 viral load (copies/ml) highlighted in bold. Blue represents positive correlation and red negative correlation. Cell types ordered by hierarchical clustering.

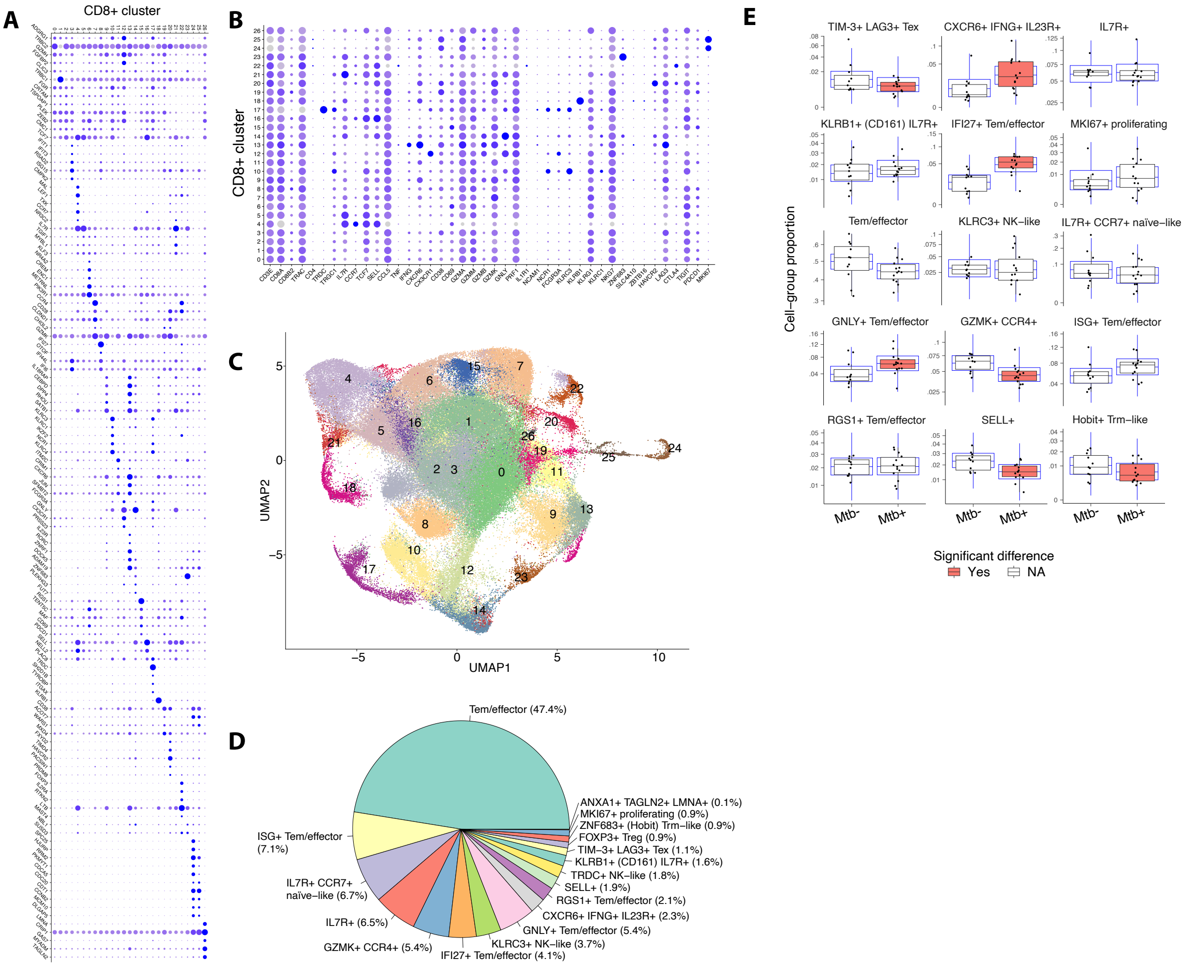


**Fig. S3. CD8 T cells**. **(A)** Dot plot of top 5 DEG per cluster. **(B)** Dot plot of canonical genes used to guide manual annotation. **(C)** Unannotated clustering of CD8+ T cells. Raw counts from 123,557 CD8A+ CD3+ cells were re-normalised, merged, integrated and clustered at resolution 0.8 to create 27 clusters. **(D)** Pie chart of CD8 T cell cluster proportions. **(E)** Comparison of CD8 T cell subcluster composition by microbiological-confirmation using the sccomp R package adjusting for age and sex. The black boxplot is the observed data; the blue boxplot is simulated data using a posterior predictive distribution. GXPU, GeneXpert Ultra. *IFNG*, interferon gamma; *GNLY*, granulysin; Tem, effector memory T cell.

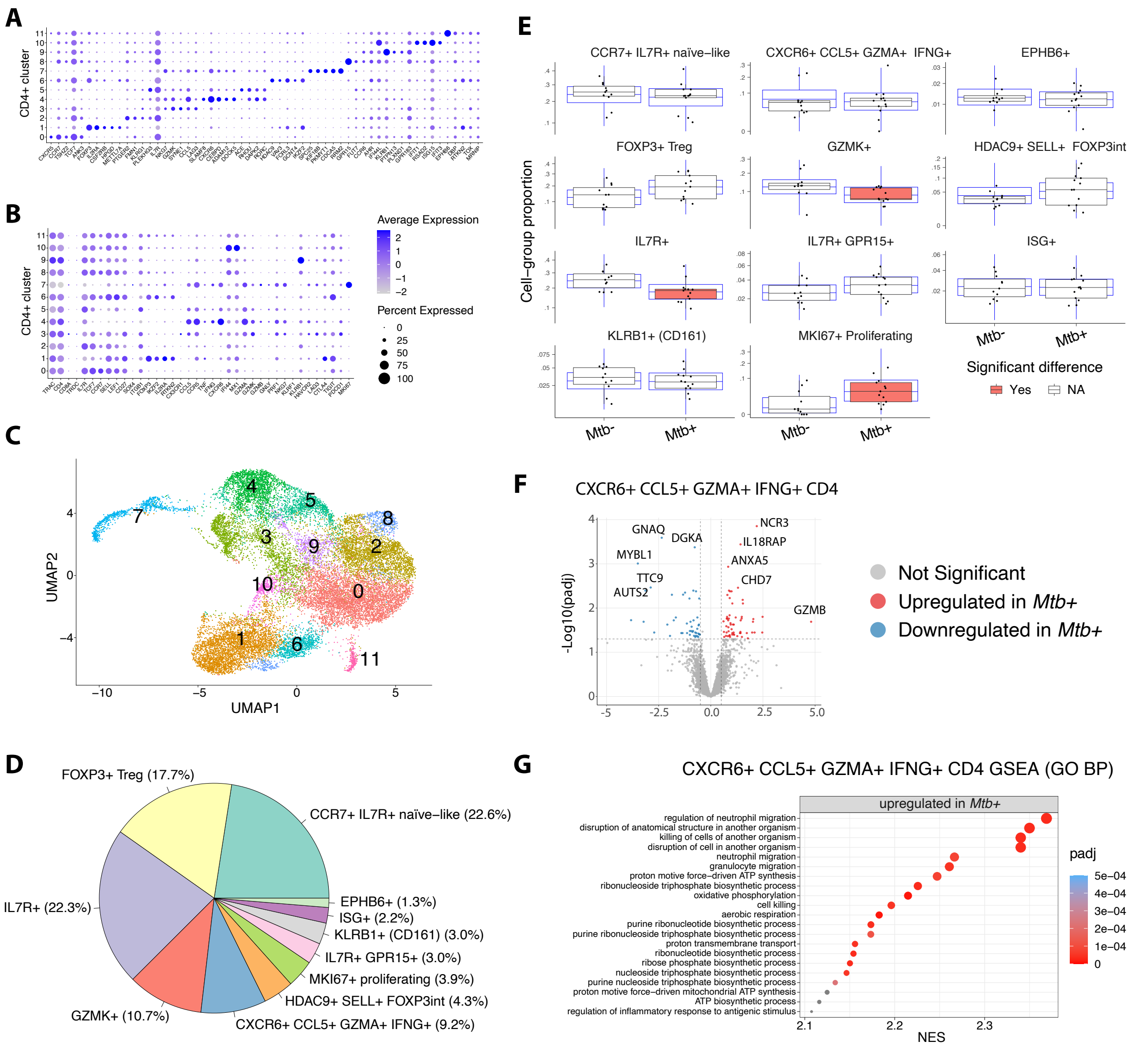


**Fig. S4. CD4 T cells**. **(A)** Dot plot of top 5 DEG per cluster. **(B)** Dot plot of canonical genes used to guide manual annotation. **(C)** Unannotated UMAP of CD4 T cells. Raw counts from 21,844 CD4+ CD3+ cells were re-normalised, merged, integrated and clustered at resolution 0.4 to create 12 clusters. **(D)** Pie chart of CD4 T cell cluster proportions. **(E)** Comparison of CD4 T cell subcluster composition by microbiological-confirmation using the sccomp R package adjusting for age and sex. The black boxplot is the observed data; the blue boxplot is simulated data using a posterior predictive distribution. **(F)** Volcano plot of DEG of CXCR6+ CCL5+ GZMA+ IFNG+ CD4 T cell by microbiological confirmation. **(G)** Gene set enrichment analysis of DEG by microbiological confirmation in the CXCR6+ CCL5+

GZMA+ IFNG+ CD4 T cell cluster done using the Gene Ontology Biological Processes gene set, ranked by log2 fold change. ISG, interferon-stimulated gene; *KLRB1*, killer cell lectin like receptor 1 (CD161); *EPHB6*, ephrin type-B receptor 6; *GZMA*, granzyme A; *IFNG*, interferon gamma.

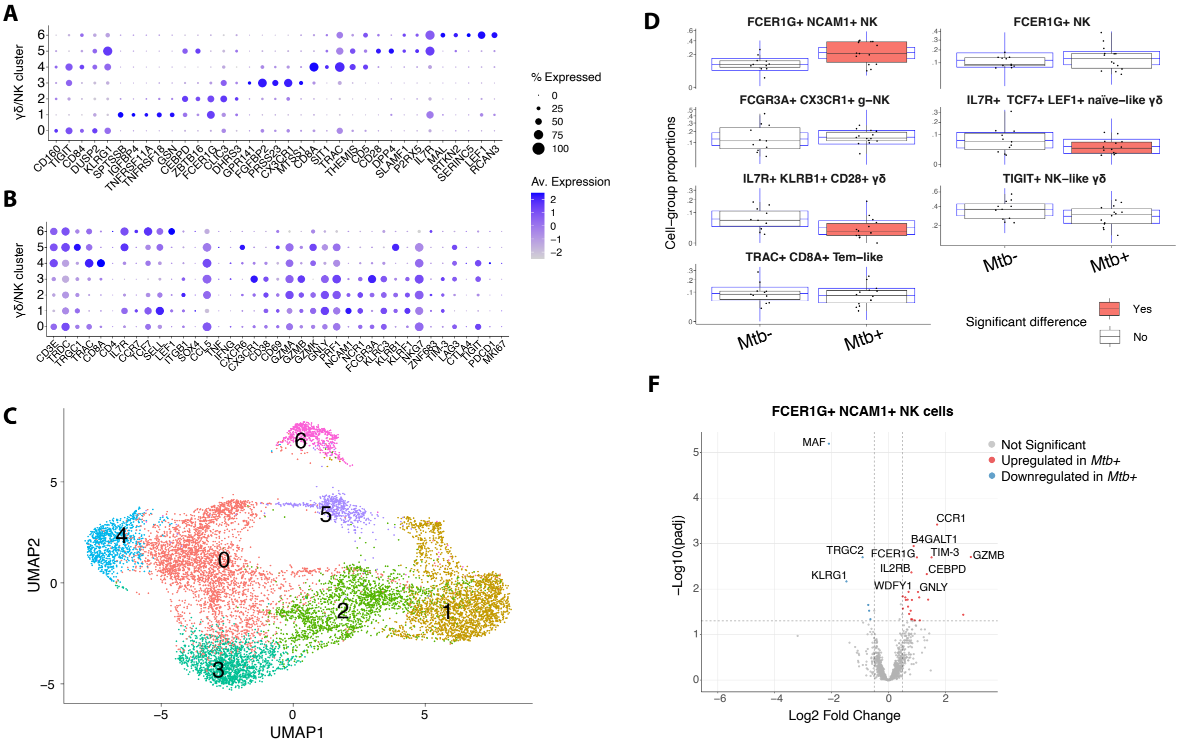


**Fig. S5. NK and γδ cells. (A)** Dot plot of top 5 DEG per cluster. **(B)** Dot plot of canonical genes used to guide manual annotation. **(C)** Unannotated UMAP of NK and γδ T cells. Raw counts from 11,483 NK and γδ cells were re-normalised, merged, integrated and clustered at resolution 0.2 to create seven clusters. **(D)** Comparison of NK and γδ T cell subcluster composition by microbiological-confirmation using the sccomp R package adjusting for age and sex. The black boxplot is the observed data; the blue boxplot is simulated data using a posterior predictive distribution. **(E)** Volcano plot of DEG for the FCER1G+ NCAM1+ NK cell cluster by microbiological confirmation. *FCER1G*, Fc fragment of IgE receptor, gamma chain; *NCAM1*, neural cell adhesion molecule 1.

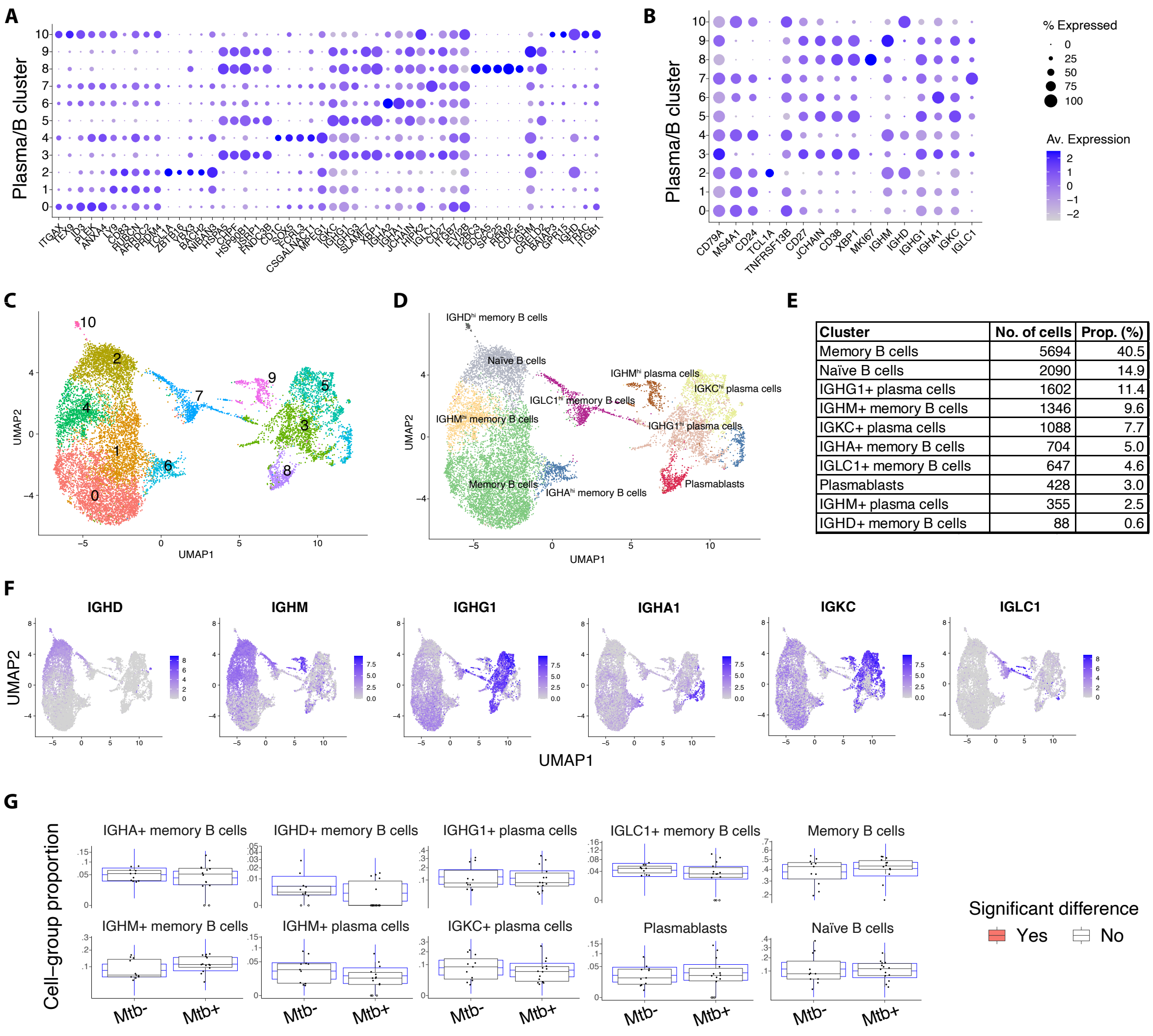


| Cluster | No. of cells | Prop. (%) |
|---|---|---|
| Memory B cells | 5694 | 40.5 |
| Naïve B cells | 2090 | 14.9 |
| IGHG1+ plasma cells | 1602 | 11.4 |
| IGHM+ memory B cells | 1346 | 9.6 |
| IGKC+ plasma cells | 1088 | 7.7 |
| IGHA+ memory B cells | 704 | 5.0 |
| IGLC1+ memory B cells | 647 | 4.6 |
| Plasmablasts | 428 | 3.0 |
| IGHM+ plasma cells | 355 | 2.5 |
| IGHD+ memory B cells | 88 | 0.6 |

**Fig. S6. B and plasma cell subclustering by microbiological confirmation. (A)** Dot plot of top 5 DEG per cluster. **(B)** Dot plot of canonical genes used to guide manual annotation. **(C)** Unannotated UMAP of B and plasma cells. Raw counts from 14,042 B cells and plasma cells were re-normalised, merged, integrated and clustered at resolution 0.3 to create 11 clusters. **(D)** Annotated UMAP of 14,042 plasma and B cells. **(E)** Table of absolute numbers and proportions of B and plasma cell clusters. **(F)** Feature plots of immunoglobulin gene expression. **(G)** Comparison of B and plasma cell subcluster composition by microbiological-confirmation using the sccomp R package adjusting for age and sex. The black boxplot is the observed data; the blue boxplot is simulated data using a posterior predictive distribution. *IGHD*, IgD heavy chain; *IGHM*, IgM heavy chain; *IGHG1*, IgG heavy chain 1; *IGHA1*, IgA heavy chain 1; *IGKC*, immunoglobulin kappa light chain constant; *IGLC1*, immunoglobin lambda light chain constant 1.

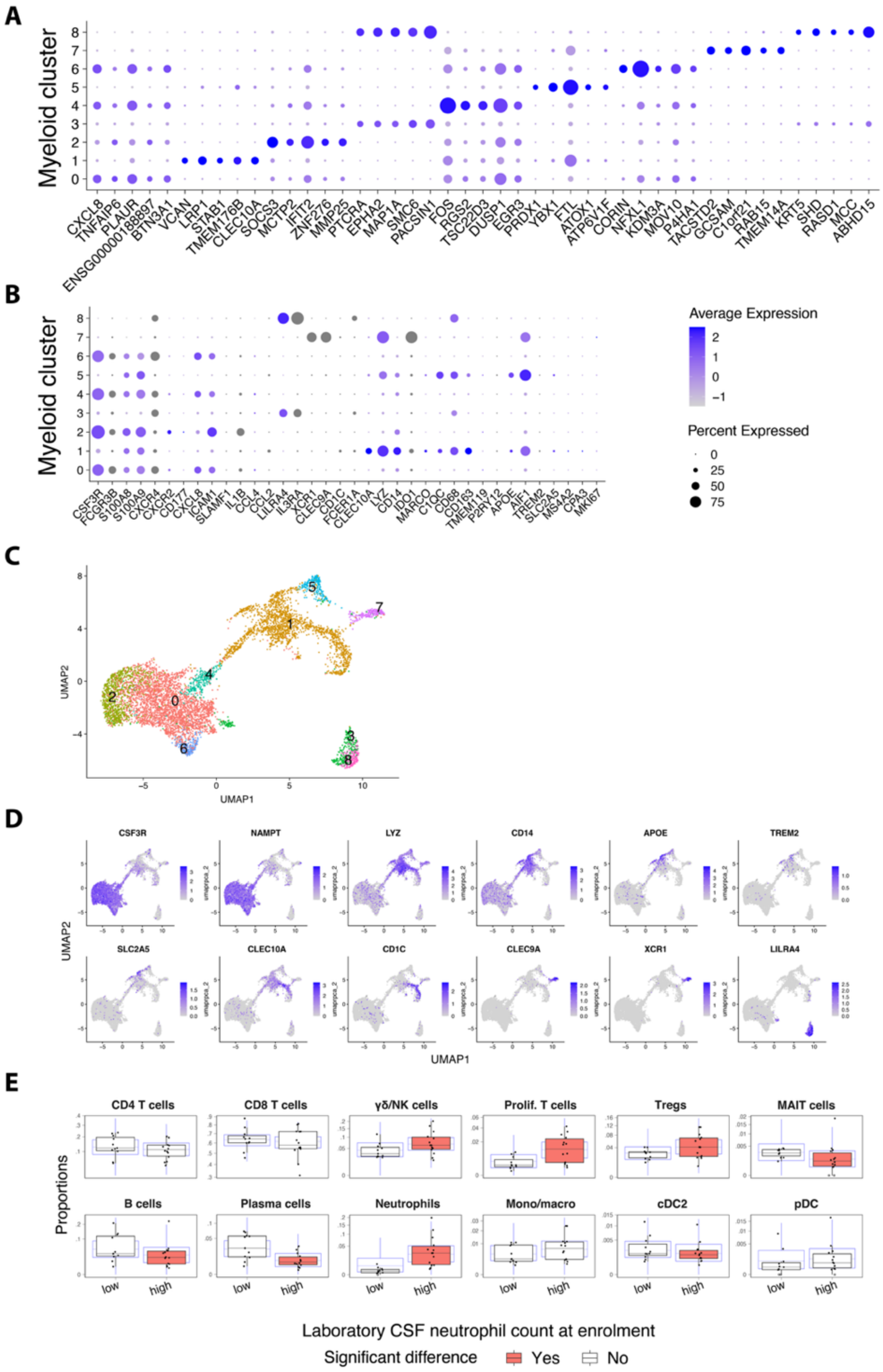
A
Myeloid cluster
B
Average Expression
Percent Expressed
C
UMAP1
UMAP2
D
CSF3R
NAMPT
LYZ
CD14
APOE
TREM2
SLC2A5
CLEC10A
CD1C
CLEC9A
XCR1
LILRA4
E
CD4 T cells
CD8 T cells
γδ/NK cells
Prolif. T cells
Tregs
MAIT cells
B cells
Plasma cells
Neutrophils
Mono/macro
cDC2
pDC
Proportions
low
high
Laboratory CSF neutrophil count at enrolment
Significant difference Yes No

**Fig. S7. Myeloid cells. (A)** Dot plot of top 5 DEG per cluster. **(B)** Dot plot of canonical genes used to guide manual annotation. **(C)** Unannotated UMAP of myeloid cells. Raw counts from 7,182 myeloid cells were re-normalised, merged, integrated and clustered at resolution 0.2 to create nine clusters. **(D)** Feature plots of myeloid cells with neutrophil canonical markers. **(E)** Comparison of coarse cell composition by laboratory CSF neutrophil count (high = ≥5 cells/mm$^3$, low = <5 cells/mm$^3$) using the sccomp R package adjusting for age and sex. The black boxplot is the observed data; the blue boxplot is simulated data using a posterior predictive distribution. *CSF3R*, colony-stimulating factor 3 receptor; *NAMPT*, nicotinamide phosphoribosyltransferase (visfatin); *LYZ*, lysozyme; *APOE*, apolipoprotein E; *TREM2*, triggering receptor expressed on myeloid cells 2; *SLC2A5*, solute carrier family 2 member 5 (GLUT5); *CLEC10A*, C-type lectin domain family 10 member A; *CLEC9A*, C-type lectin domain family 9 member A; *XCR1*, XC motif chemokine receptor 1; *LILRA4*, leukocyte immunoglobulin-like receptor subfamily A member 4.

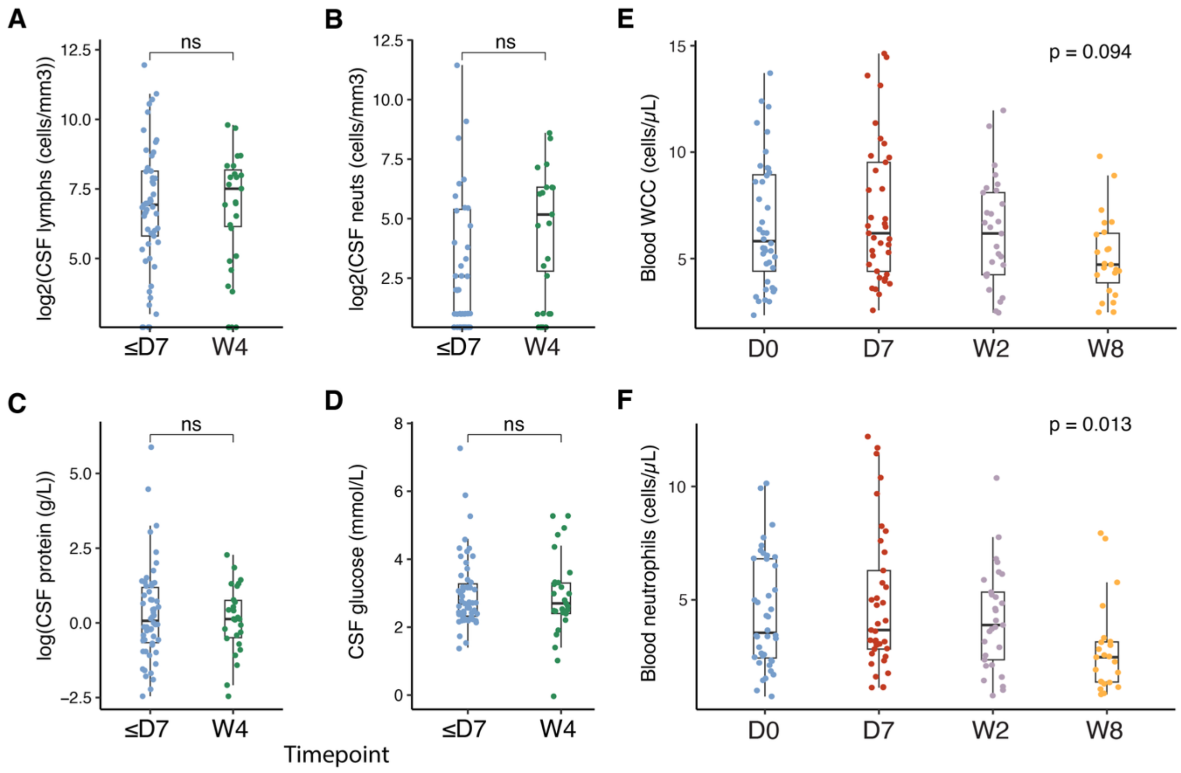


**Fig. S8. Cell counts over time for participants in CSF and blood bulk RNA sequencing**.

Comparison between early (≤7 days, n = 44) and late (week 4, n = 20) laboratory lymphocyte counts (A), neutrophil counts (B), protein (C) and glucose (D) for participants enrolled in INTENSE-TBM and LASER-TBM whose samples were used for CSF bulk RNA sequencing (patients = 53, samples = 64). +1 added to CSF cell counts, and +0.1 added to CSF protein to allow for log2 transformation of zero values. Comparisons between laboratory white cell count (E) and neutrophil count (F) at day 0 (n = 40), day 7 (n = 35), week 2 (n = 28) and week 8 (n = 22) for participants enrolled in LASER-TBM whose samples were used for blood bulk RNA sequencing (patients = 42, samples = 125). Comparisons made using Mann Whitney U to compare the two CSF timepoints and ANOVA to compare the four blood timepoints. ns = non-significant.

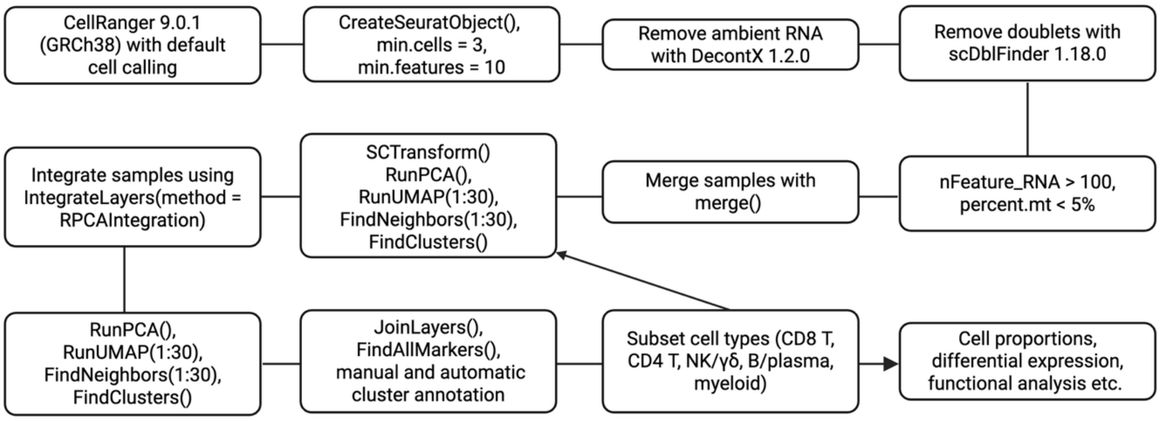


**Fig. S9. 10x Flex single cell RNA sequencing bioinformatics workflow.** Doublets refer to two or more cells that have been captured in the same droplet at the cell separation step, and therefore share the same cellular barcode. SCTransform refers to a normalisation and variance stabilisation technique using regularised negative binomial regression. RPCA, reciprocal principal component analysis.

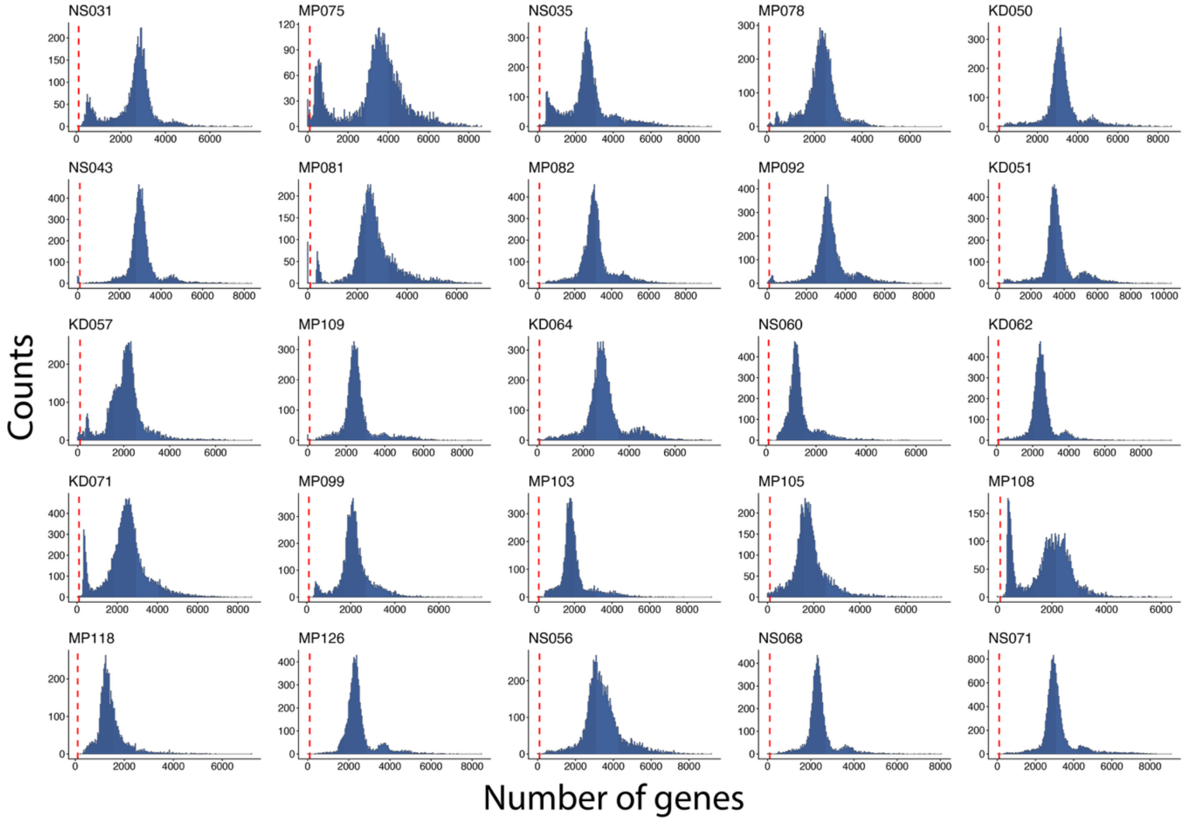


**Fig. S10. Histograms of genes per cell**. Histogram of each sample plotting the number of genes per cell (nFeature_RNA) vs. frequency. Red dotted line indicates the cut off of 100 genes/cell used for filtering. Each panel is one donor.

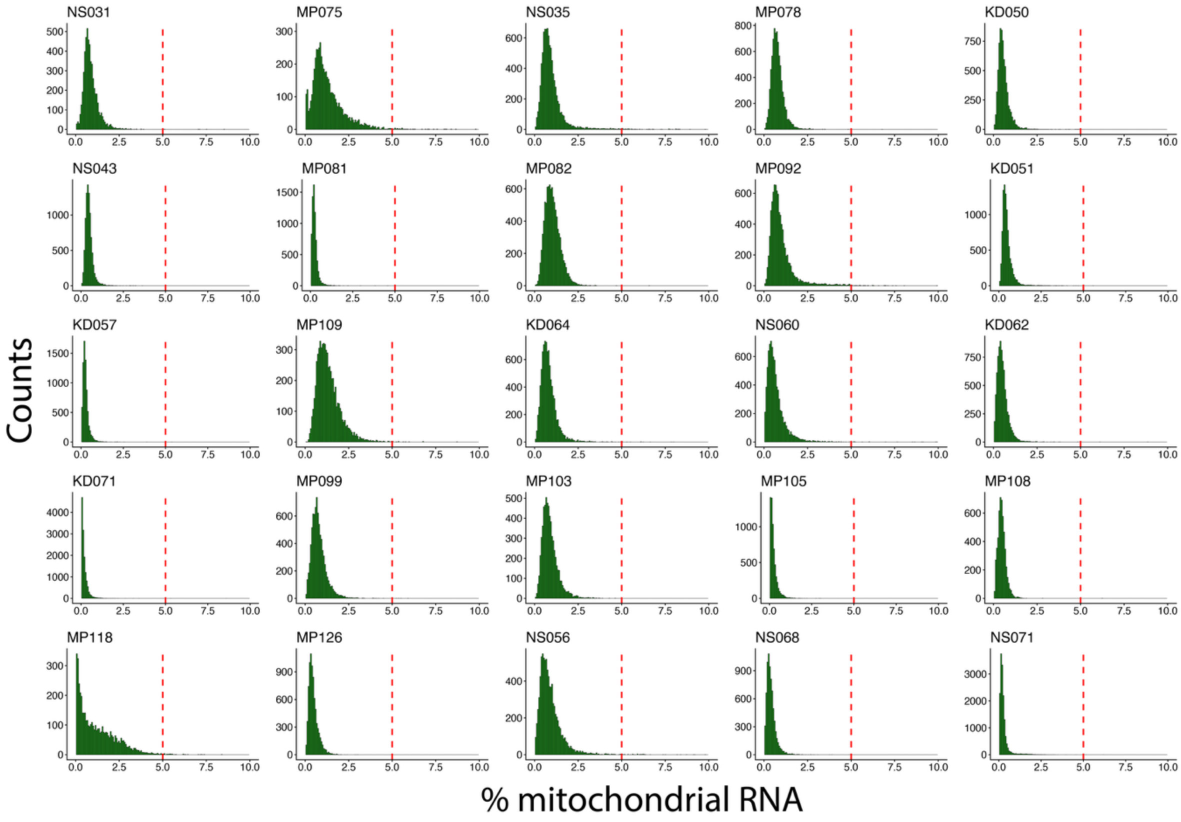


**Fig. S11. Histograms of percentage of mitochondrial RNA per cell**. Histogram of each sample showing the percentage of mitochondrial RNA per cell vs. frequency. Red dotted line indicates the cut off of 5% mitochondrial RNA used for filtering. Each panel is one donor.

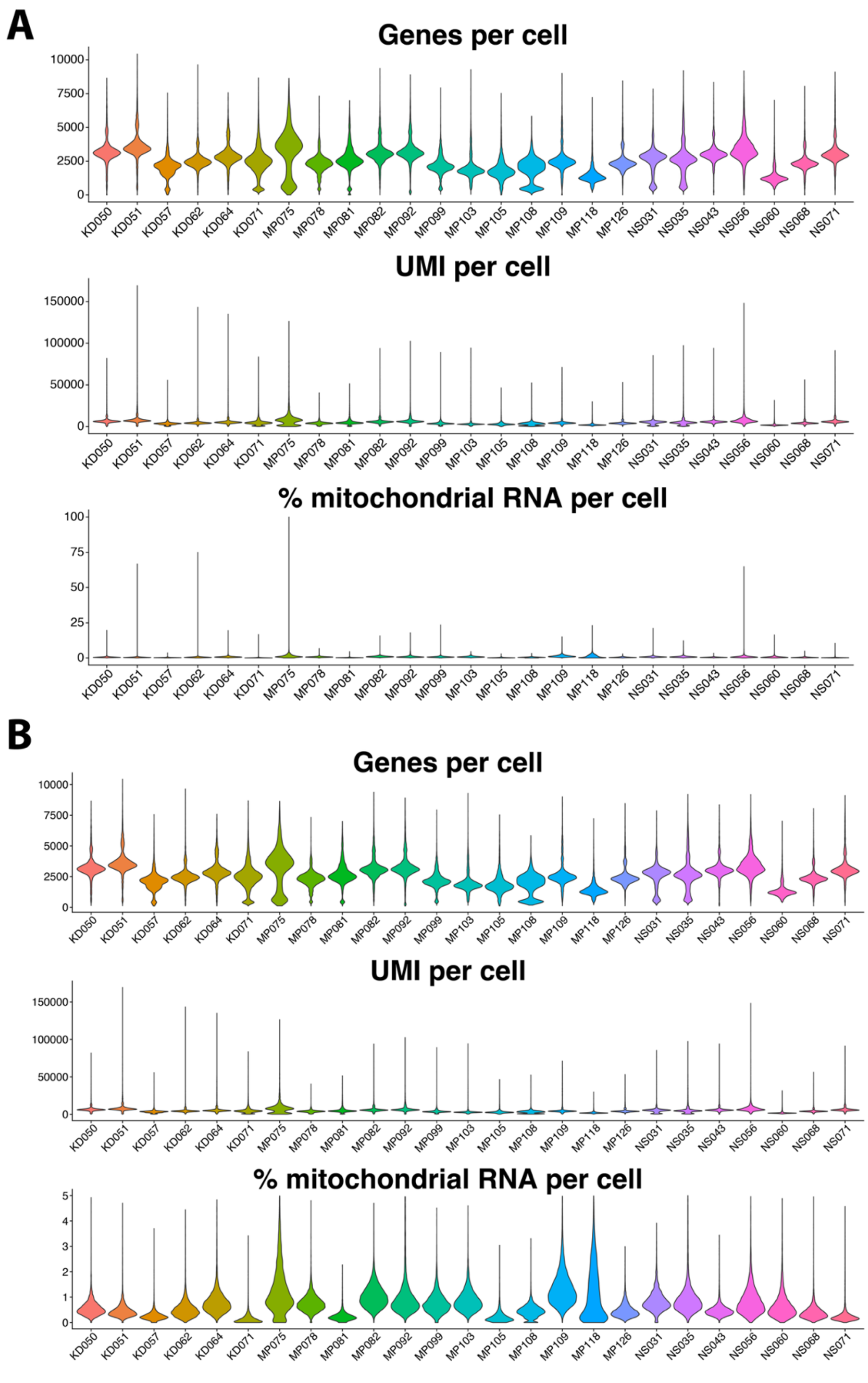


**Fig. S12. Genes, UMI and percent mitochondrial genes per cell before and after filtering**. A) Genes, UMI and percent mitochondrial genes per cell before filtering B) Genes, UMI and percent mitochondrial genes per cell after filtering cells with <100 genes and >5% mitochondrial RNA.

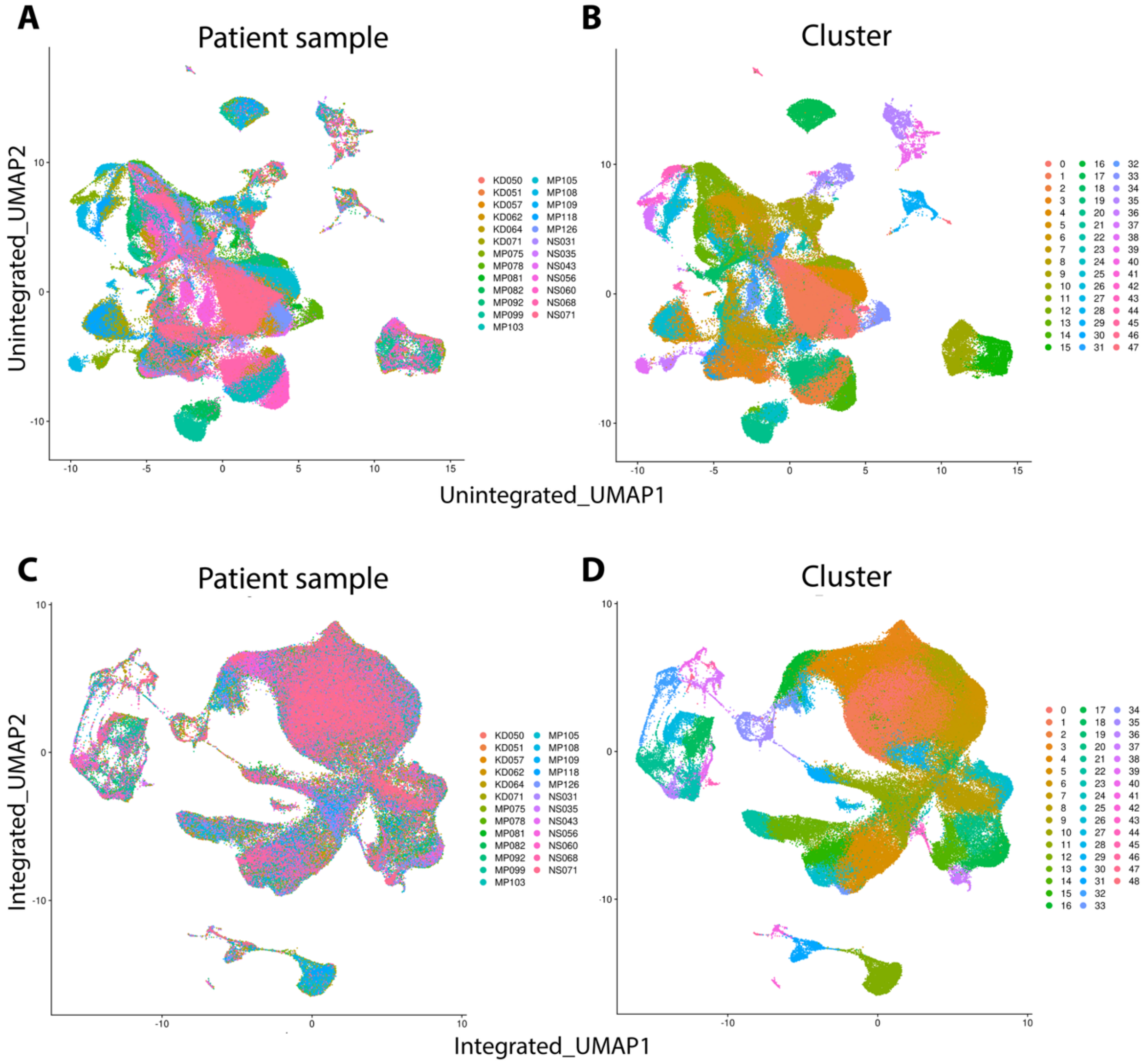


**Fig. S13. Reciprocal PCA integration. (A)** Unintegrated UMAP coloured by sample. **(B)** Unintegrated UMAP coloured by default clustering. **(C)** Reciprocal PCA integrated UMAP coloured by sample. **(D)** Reciprocal PCA integrated UMAP coloured by clustering at resolution 0.9. N.B. Sample colours are added by layer, with NS071 added last, meaning that later samples appear more prominent.

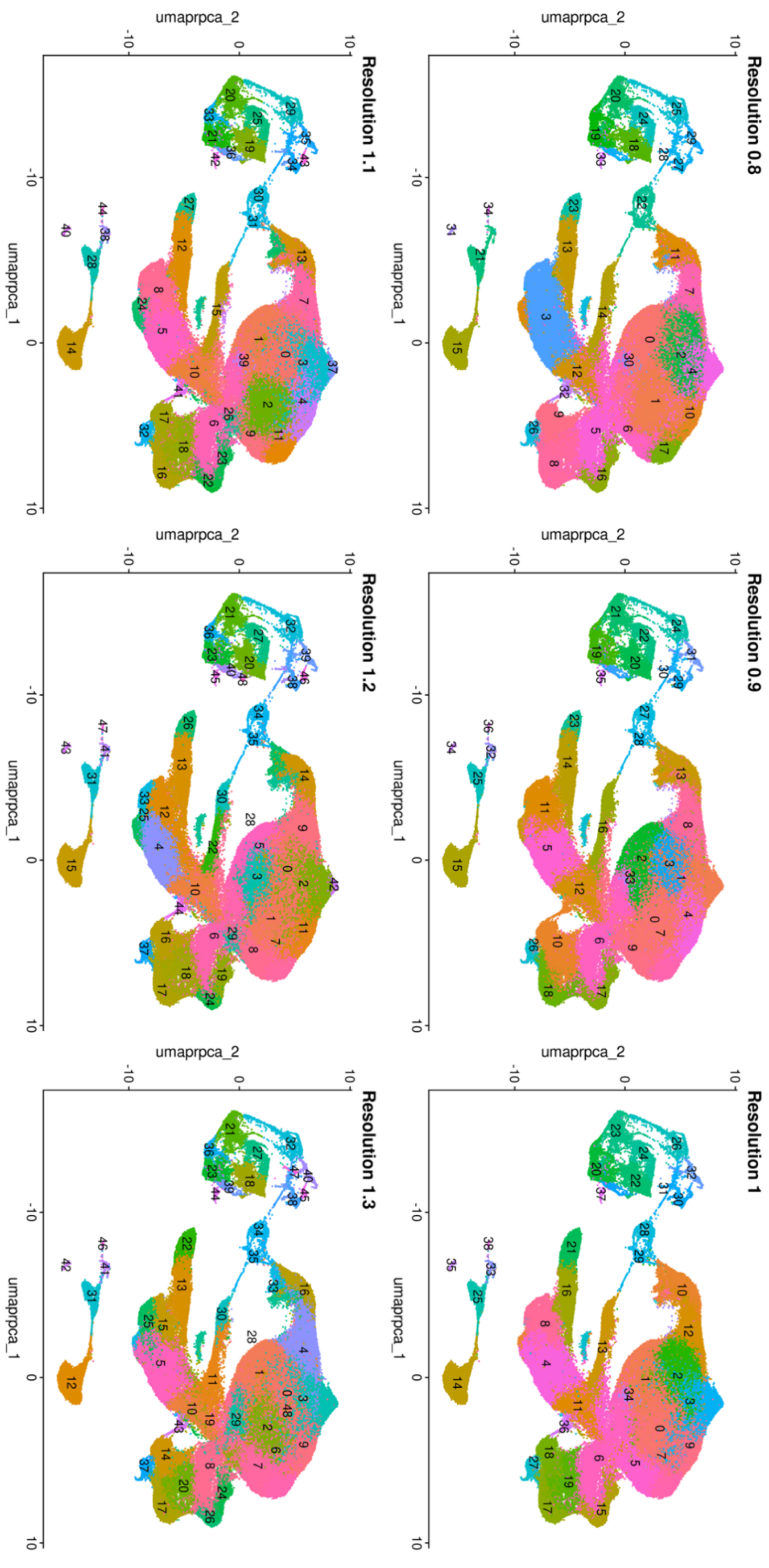


**Fig. S14. Integrated UMAPs at selected cluster resolutions**. UMAP at six resolutions using FindClusters(). Resolution 0.9 was used for annotation.

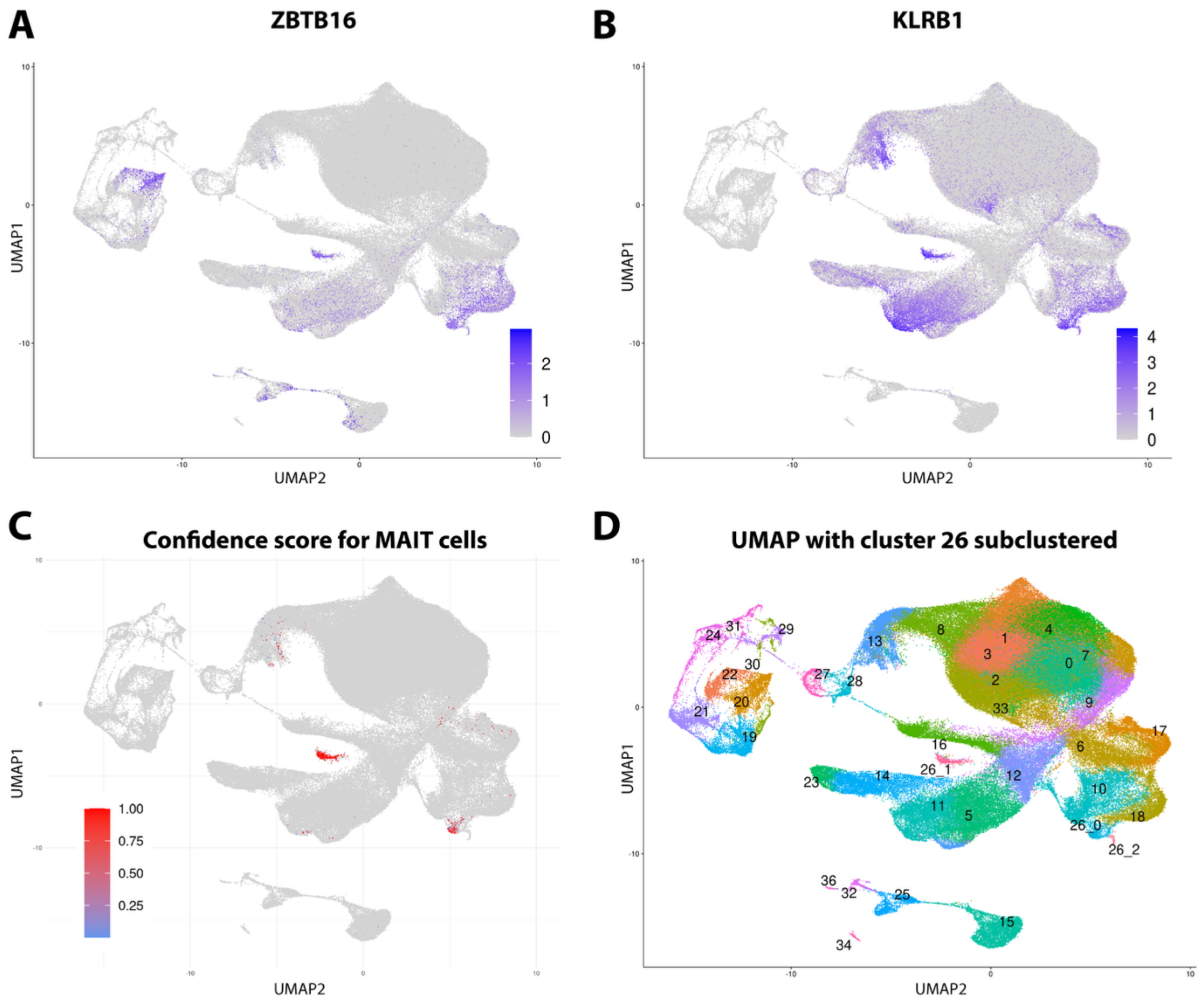


**Fig. S15. Annotation of MAIT cells. (A)** UMAP highlighting *ZBTB16* (zinc finger and BTB domain-containing protein 16, promyelocytic leukaemia zinc finger, PLZF) gene expression. **(B)** UMAP highlighting *KLRB1* (killer cell lectin-like receptor subfamily B, member 1, CD161). **(C)** Celltypist confidence scores for cells labelled as MAIT cells in data. Red means high confidence, blue means low confidence. **(D)** UMAP of all cells after subclustering cluster 26 using FindSubClusters() into three smaller clusters. Cluster 26_1 (n = 501) was re-labelled as MAIT cells based on the Celltypist confidence and canonical gene expression.
